# Myosin-driven Nucleation of Actin Filaments Drives Stereocilia Development Critical for Hearing

**DOI:** 10.1101/2021.07.09.451618

**Authors:** Zane G. Moreland, Fangfang Jiang, Carlos Aguilar, Melanie Barzik, Rui Gong, Arik Shams, Christian Faaborg-Andersen, Jesse C. Werth, Randall Harley, Daniel C. Sutton, Stacey M. Cole, Andrew Parker, Susan Morse, Elizabeth Wilson, Yasuharu Takagi, James R. Sellers, Steve D.M. Brown, Thomas B. Friedman, Gregory M. Alushin, Michael R. Bowl, Jonathan E. Bird

## Abstract

The assembly and maintenance of actin-based mechanosensitive stereocilia in the cochlea is critical for lifelong hearing. Myosin-15 (MYO15) is hypothesized to modulate stereocilia height by trafficking actin regulatory proteins to their tip compartments, where actin polymerization must be precisely controlled during development. We identified a mutation (p.D1647G) in the MYO15 motor-domain that initially maintained trafficking, but caused progressive hearing loss by stunting stereocilia growth, revealing an additional function for MYO15. Consistent with its maintenance of tip trafficking *in vivo*, purified p.D1647G MYO15 modestly reduced actin-stimulated ATPase activity *in vitro*. Using ensemble and single-filament fluorescence *in vitro* assays, we demonstrated that wild-type MYO15 directly accelerated actin filament polymerization by driving nucleation, whilst p.D1647G MYO15 blocked this activity. Collectively, our studies suggest direct actin nucleation by MYO15 at the stereocilia tip is necessary for elongation *in vivo*, and that this is a primary mechanism disrupted in DFNB3 hereditary human hearing loss.

## INTRODUCTION

Cochlear hair cells are the primary transducers of sound in the mammalian inner ear and are fundamental for hearing. Each hair cell assembles approximately 100 individual stereocilia on its apical surface to form a mechano-sensitive hair bundle. Within each bundle, stereocilia are precisely graded into ranks of ascending height and this staircase architecture is critical for mechano-electric transduction (MET) (Barr-Gillespie, 2015). The regulation of stereocilia size is thus central for hearing and the disruption of hair bundle architecture is a common theme in hereditary deafness (Richardson et al., 2011). Stereocilia develop from microvilli by building a highly cross-linked para-crystalline core of actin filaments as an internal scaffold to confer shape and structural rigidity (Vélez-Ortega and Frolenkov, 2019). As stereocilia develop, the actin core thickens and elongates to reach its mature size, necessitating precise control of actin filament polymerization (Tilney et al., 1992; Kaltenbach et al., 1994; Krey et al., 2020). Actin filaments are uniformly polarized within stereocilia with rapidly growing barbed ends orientated towards the tip, the major site of actin polymerization and filament elongation, and also the site of MET (Flock and Cheung, 1977; Tilney et al., 1980; Beurg et al., 2009). Once stereocilia are fully assembled, actin polymerization continues primarily at the tip compartment, indicating an ongoing plasticity throughout adult life (Schneider et al., 2002; Zhang et al., 2012; Drummond et al., 2015; Narayanan et al., 2015). A number of proteins have been identified within the tip compartment that are essential for stereocilia growth, yet the molecular mechanisms governing actin polymerization are unknown. Identification of this mechanism is central to the acquisition and maintenance of hair cell mechano-sensitivity, and also for understanding hearing loss as the clinical manifestation of a cytoskeletal disease.

A key molecule in establishing stereocilia architecture is unconventional myosin 15 (MYO15), encoded by the gene *Myo15* in mice and *MYO15A* in humans. Mutations in *MYO15A* (MIM #602666) cause DFNB3 recessive hereditary hearing loss in humans (Friedman et al., 1995; Wang et al., 1998; Rehman et al., 2016). MYO15 is a member of the myosin superfamily of P-loop ATPases that generate contractile force on actin filaments to power cellular processes such as cytokinesis, endocytosis and vesicular trafficking (Hartman and Spudich, 2012; Houdusse and Sweeney, 2016). Two MYO15 isoforms are produced in auditory hair cells through alternative mRNA splicing (Liang et al., 1999; Anderson et al., 2000; Fang et al., 2015). A shorter isoform (MYO15-2, also known as MYO15-S) consists of the ATPase ‘motor domain’ and three light chain binding sites that associate with calmodulin-like proteins, in addition to myosin tail homology 4 (MyTH4), Src homology 3 (SH3), and protein 4.1, ezrin, radixin, moesin (FERM) domains (Fig. 1A). A larger isoform (MYO15-1, also known as MYO15-L) is identical to MYO15-2, except for the addition of a 133 kDa N-terminal domain (Fig. 1A). Both MYO15 isoforms localize to the stereocilia tip compartment where actin polymerization is concentrated (Schneider et al., 2002; Belyantseva et al., 2003; Drummond et al., 2015; Fang et al., 2015; Narayanan et al., 2015). The prototypical *Myo15* mutant allele, the *shaker 2* (*Myo15^sh2^*), prevents both isoforms from accumulating in stereocilia, blocking developmental elongation and causing profound hearing loss from birth (Probst et al., 1998; Anderson et al., 2000; Belyantseva et al., 2003; Fang et al., 2015). The use of an isoform-specific *Myo15* null allele revealed additional functions, with MYO15-2 being necessary for stereocilia developmental elongation, whilst MYO15-1 independently maintains the postnatal size of shorter stereocilia (Fang et al., 2015) that are structurally plastic in response to MET (Vélez-Ortega et al., 2017; Krey et al., 2020).

**FIGURE 1.**
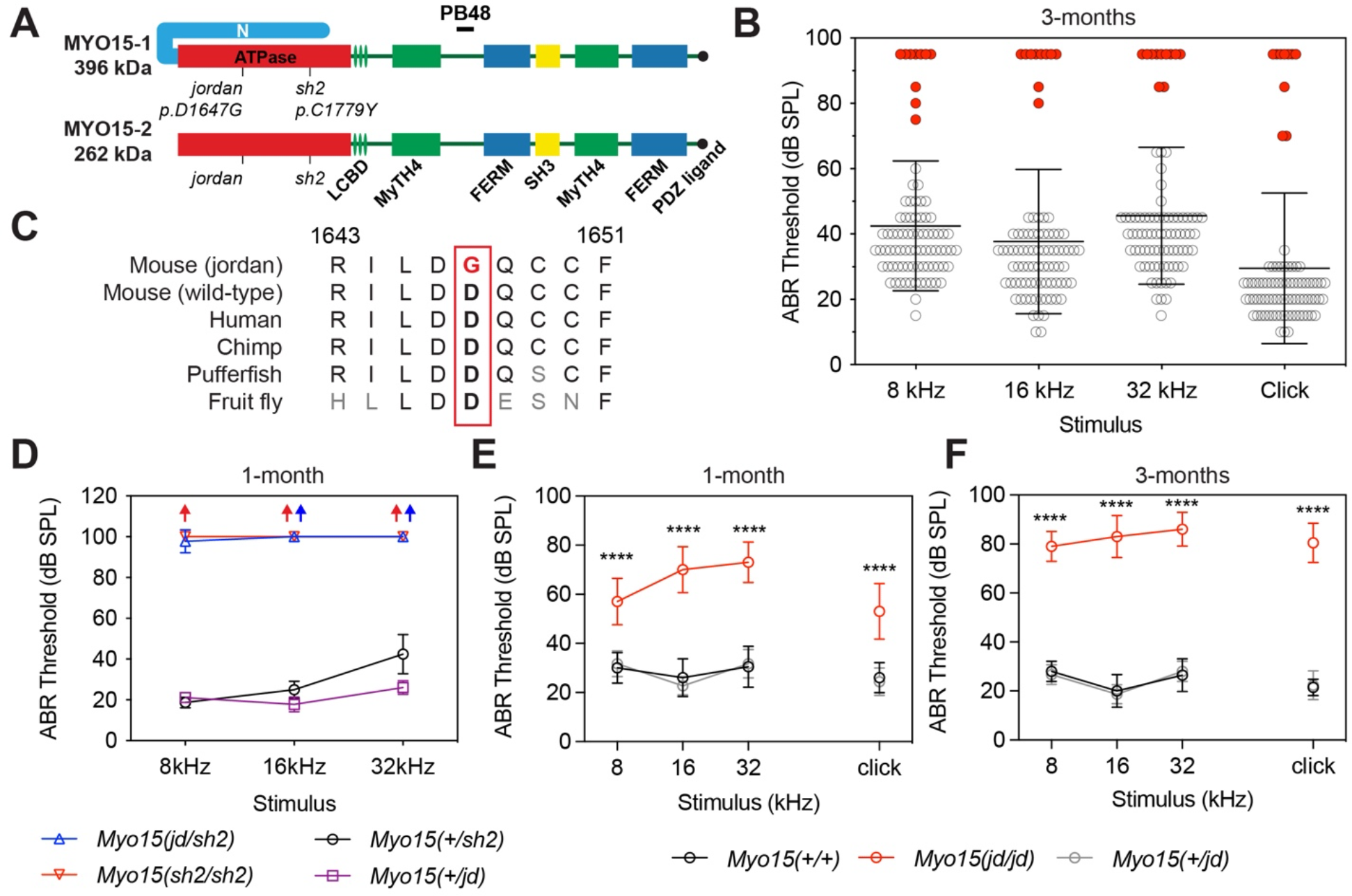
**A)** Schematic showing the protein domains of the long (MYO15-1) and short (MYO15-2) isoforms encoded by the *Myo15* gene. The *jordan* and *shaker-2* deafness mutations are shown. **B)** ABR phenotyping of the *jordan* pedigree at 3 months identified 10 mice with statistically elevated hearing thresholds (red circles) for click and at 8, 16 and 32 kHz stimuli, compared to their normal hearing pedigree mates (n = 73, black circles). Statistical outliers were detected using robust regression and outlier removal (red circles, ROUT, Q = 1%). Thresholds of affected mice that did not respond to the highest intensity stimulus (90 dB SPL) are recorded as 95 dB SPL. Data are mean ± SD. **C)** Evolutionary conservation of the aspartate (D) residue of MYO15 altered to glycine (G) in *jordan* mice that causes hearing loss. Residue positions refer to mouse MYO15-1 (NP_034992.2). **D)** ABR recordings of *Myo15^jd/sh2^* compound heterozygotes at P28 shows profound hearing loss, with thresholds elevated compared with normal hearing *Myo15^+/jd^ or Myo15^+/sh2^* littermates. **E-F)** Longitudinal auditory phenotyping of *jordan* mice at 1-(E) and 3-(F) months of age. ABR recordings show that *Myo15^jd/jd^* mice (red, n=10) exhibit a progressive, moderate-to-severe hearing loss affecting all frequencies, whereas age-matched *Myo15^+/+^*(black, n=10) and *Myo15^+/jd^* (grey, n=15) littermate controls have normal thresholds (15 - 45 dB SPL). *Myo15^+/+^ vs Myo15^jd/jd^* comparison, ANOVA with Tukey’s multiple comparisons test.

The MYO15-2 isoform associates with additional proteins essential for stereocilia elongation, and by inference, actin polymerization. The ‘elongation complex’ consists of epidermal growth factor receptor pathway substrate 8 (EPS8), whirlin (WHRN), G-protein signalling modulator 2 (GPSM2) and G-protein subunit alphai3 (GNAI3). Similar to MYO15-2, these proteins are concentrated at the tips of the tallest stereocilia (row 1) (Mburu et al., 2003; Belyantseva et al., 2005; Delprat et al., 2005; Manor et al., 2011; Zampini et al., 2011; Tarchini et al., 2016; Mauriac et al., 2017; Tadenev et al., 2019). Individual knock-out mouse mutants of *Eps8*, *Whrn*, *Gpsm2* or *Gnai3* recapitulate the *shaker 2* phenotype, exhibit short stereocilia and are profoundly deaf (Mburu et al., 2003; Manor et al., 2011; Zampini et al., 2011; Tarchini et al., 2016; Mauriac et al., 2017; Tadenev et al., 2019). There is strong evidence that a key function of MYO15-2 is to traffic the elongation complex and concentrate it at the stereocilia tips. First, elongation complex proteins are absent from the stereocilia of *Myo15 shaker 2* hair cells, demonstrating their functional dependence upon MYO15-2 *in vivo* (Belyantseva et al., 2005; Delprat et al., 2005; Manor et al., 2011; Mauriac et al., 2017; Tadenev et al., 2019). Second, EGFP-tagged MYO15-2 actively traffics elongation complex proteins along filopodia in cell lines (Belyantseva et al., 2005; Manor et al., 2011; Mauriac et al., 2017). Finally, enzymatic studies of the purified MYO15 ATPase domain reveal kinetic adaptations that enable long-range processive molecular trafficking (Bird et al., 2014; Jiang et al., 2021). Together, these data support MYO15-2 delivering the elongation complex to the stereocilia tips, where the elongation complex is hypothesized to regulate actin polymerization. Despite its central role in promoting stereocilia growth, the specific molecular activity of the elongation complex remains unknown.

Here, we describe a novel *Myo15* mutant mouse (‘*jordan*’) that exhibits progressive hearing loss resulting from a missense substitution in the MYO15 motor domain. In striking contrast with *shaker 2* hair cells, MYO15 and the elongation complex proteins are still delivered to the stereocilia tip compartment in *jordan* mutant hair cells, but stereocilia fail to elongate properly. These results questioned the sufficiency of the elongation complex to drive stereocilia growth, and prompted us to search for another role of MYO15 independent of delivering the elongation complex. We found that purified MYO15 motor domain protein directly stimulated actin polymerization *in vitro*, and that the *jordan* mutation blocked this activity, whilst only moderately affecting its ability to bind and move along actin filaments. A companion structural study shows that the *jordan* mutation targets the actomyosin binding interface and interferes with the ability of wild-type MYO15 to regulate structural plasticity within the actin molecule itself (Gong et al., 2021). Combined, our results argue that MYO15 controls stereocilia elongation by directly regulating F-actin conformation in stereocilia, and that this critical activity is independent of the elongation complex. More broadly, our work suggests that in addition to their classical roles generating force and motility, myosin motors have a physiological role regulating actin polymerization *in vivo*.

## RESULTS

### A forward genetic screen identifies *jordan,* a *Myo15* allele causing progressive hearing loss

During a recent phenotype-driven ENU-mutagenesis screen (Potter et al., 2016), the MPC190 cohort (comprising 83 mice) was identified with 10 mice exhibiting severe hearing loss at 3-months of age (Fig. 1B). A genome scan and single nucleotide polymorphism (SNP) mapping of third generation (G3) deaf mice found linkage to a 16.7 Mb region on Chromosome 11 (Fig. S1A). Whole-genome sequencing of a single deaf mouse identified a high confidence homozygous mutation within the critical interval, consisting of an A-to-G transition at coding nucleotide 4940 of the *Myo15* gene (ENSMUST00000071880). This variant was confirmed by Sanger sequencing (Fig. S1B) and leads to the substitution of an evolutionarily conserved aspartate residue with a glycine (p.D1647G) in the encoded MYO15 protein (Fig. 1C). We named this mutant allele *jordan* (*Myo15^jd^*).

To confirm that the *Myo15^jd^* substitution causes hearing loss, we performed a genetic complementation test utilizing the *shaker-2* (*Myo15^sh2^*) deafness allele *in trans* (Probst et al., 1998; Stepanyan et al., 2006). We recorded ABRs from postnatal day 28 (P28) mice and found that compound heterozygous *Myo15^jd/sh2^* mice had elevated thresholds of >90 decibel sound pressure level (dB SPL) at all frequencies (Fig. 1D). In contrast, *Myo15^jd/+^* and *Myo15^sh2/+^* littermates had normal thresholds (< 40 dB SPL) (Fig. 1D). Failure of complementation in *Myo15^jd/sh2^* mice confirms that the p.D1647G mutation in *Myo15* is the cause of recessive deafness in the *jordan* pedigree.

All reported mutant *Myo15* mouse alleles cause profound deafness (MGI:1261811), measured from P14 onwards (Probst et al., 1998; Anderson et al., 2000; Stepanyan et al., 2006; Fang et al., 2015). ABR showed that several *Myo15^jd/jd^* mice had residual hearing at 3 months (Fig. 1B, red circles), suggesting a distinct mechanism of hearing loss. We investigated this using longitudinal ABR measurements. At 4 weeks of age, *Myo15^jd/jd^* mice had moderate hearing loss with broadband click ABR threshold of 53 ± 11 dB SPL, compared to normal hearing *Myo15^+/+^* (26 ± 6 dB SPL) and *Myo15^+/jd^* (24 ± 6 dB SPL) littermates (Fig. 1E). The hearing of *Myo15^jd/jd^* mice progressively worsened, with click ABR thresholds of 53 ± 11, 69 ± 9, 80 ± 9 and 81 ± 8 dB SPL at 4, 6, 9 and 12 weeks, respectively (Fig. 1E,F, Fig. S1C-F). Click ABR thresholds for control *Myo15^+/+^* and *Myo15^jd/+^* littermates ranged between 22 ± 4 and 26 ± 6 dB SPL; showing they did not develop hearing loss with age (Fig. 1E,F, Fig. S1C-F). To investigate outer hair cell (OHC) function we measured distortion-product otoacoustic emissions (DPOAEs), and found they were absent in *Myo15^jd/jd^* mice at 12 weeks, except for frequencies < 10 kHz, where they were significantly reduced compared to *Myo15^+/+^* and *Myo15^+/jd^* littermates (Fig. S1G). The absence of DPOAEs shows that OHC function is impaired in *Myo15^jd/jd^* mice. As the only known mouse *Myo15* variant to cause progressive hearing loss, the *jordan* allele is an important new model to explore the full spectrum of DFNB3 deafness, which presents heterogeneously as either profound congenital, or progressive hearing loss in humans (Rehman et al., 2016).

### Stereocilia do not properly elongate in *jordan* mutant hair cells

To investigate the cellular pathology underlying hearing loss in *Myo15^jd/jd^* mice, we used scanning electron microscopy (SEM) to assess the structure of cochlear hair bundles. In wild-type *Myo15^+/+^* IHCs and OHCs at P8, stereocilia were graded into a staircase pattern with three distinct rows (Fig. 2A,B). By comparison, bundle heights were reduced in *Myo15^jd/jd^* littermates, whilst still retaining the staircase architecture (Fig. 2C,D). Quantification of row 1 stereocilia heights showed that *Myo15^jd/jd^* IHCs and OHCs were significantly shorter than *Myo15^+/+^* controls (Fig. 2G,H). Furthermore, stereocilia at the lateral edge of *Myo15^jd/jd^* OHCs, and to lesser extent IHCs, were shorter in height such that the central stereocilia were tallest (Fig. 2C,D). *Myo15^jd/jd^* hair cells consistently had 1 or 2 additional rows of stereocilia (Fig. 2C,D) when compared to the three well-defined rows in *Myo15^+/+^* hair cells (Fig. 2A,B). The additional row phenotype of *Myo15^jd/jd^* hair cells was similar to *Myo15^sh2/sh2^* hair cells at P8 (Fig. 2E,F), however *Myo15^jd/jd^* stereocilia lengths were significantly longer when compared to *Myo15^sh2/sh2^* hair cells (Fig. 2G,H). We conclude that stereocilia in *Myo15^jd/jd^* hair cells elongate more than *Myo15^sh2/sh2^* hair cells, but fail to reach normal wild-type heights. The *Myo15 shaker 2* and *jordan* allelic series shows that mutations in the motor domain led to altered stereocilia heights and explain why *jordan* mice initially have less severe hearing loss than *shaker-2* mice.

**FIGURE 2.**
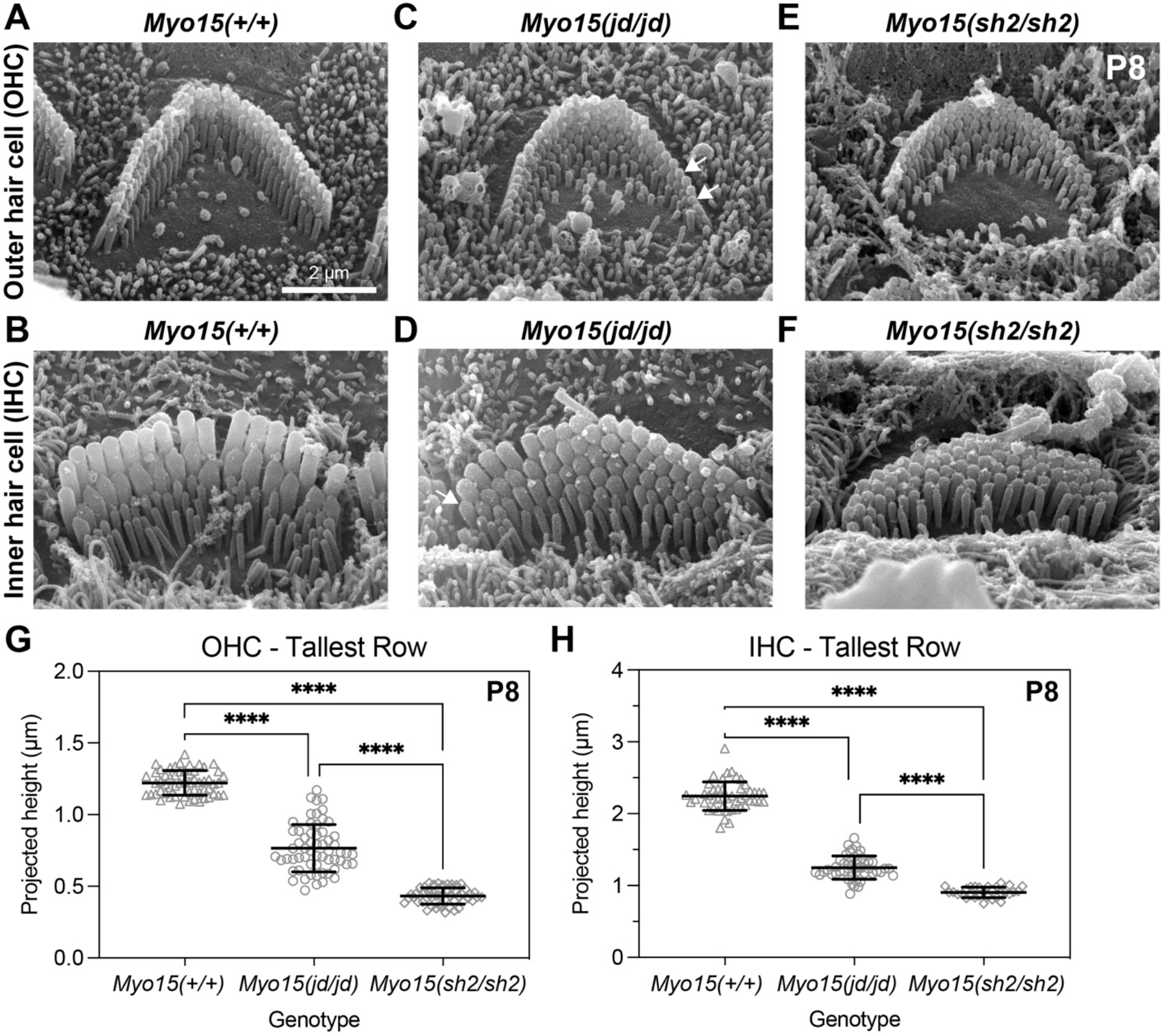
**A-F)** Representative SEM images of stereocilia bundles from *Myo15^+/+^*, *Myo15^jd/jd^*, and *Myo15^sh2/sh2^* OHCs (A,C,E) and IHCs (B,D,F) at P8. In *Myo15^+/+^* mice, both IHC and OHC bundles display the characteristic staircase architecture with 3 stereocilia ranks of increasing height. IHC and OHC bundles from either *Myo15^jd/jd^* and *Myo15^sh2/sh2^* mice are shorter in height than the wild-type bundles. *Myo15^jd/jd^* stereocilia taper in height towards the periphery of the bundle (white arrows). **G)** Projected heights of tallest (row 1) OHC stereocilia at P8 are 1.2 ± 0.1 µm (*Myo15^+/+^*, 58 stereocilia from 4 mice), 0.8 ± 0.1 µm (*Myo15^jd/jd^*, 60 stereocilia from 4 mice), and 0.4 ± 0.1 µm (*Myo15^sh2/sh2^,* 50 stereocilia from 2 mice). **H)** Projected heights of tallest (row 1) IHC stereocilia at P8 are 2.2 ± 0.2 µm (*Myo15^+/+^,* 47 stereocilia from 4 mice), 1.3 ± 0.1 µm (*Myo15^jd/jd^*, 47 stereocilia from 4 mice), and 0.9 ± 0.1 µm (*Myo15^sh2/sh2^,* 30 stereocilia from 2 mice). Brown-Forsythe and Welch ANOVA with Dunnett’s T3 multiple comparisons test. Images are from mid-cochlear turn. Scale bars, 1 µm.

To further investigate the progressive component of hearing loss, we next examined hair bundle morphology at 12 weeks, when *jordan* mice were profoundly deaf (Fig. 1F). Compared with *Myo15^+/+^*, IHC bundles of *Myo15^jd/jd^* mice had lost their staircase architecture (Fig. S2A) and were still significantly shorter (Fig. S2B). Since the staircase was initially present at P8 in *Myo15^jd/jd^* IHCs (Fig. 2D), these data suggested a postnatal resorption of stereocilia. Consistent with this hypothesis, extremely short stereocilia at the hair bundle periphery were observed at 12 weeks in *Myo15^jd/jd^* OHCs (Fig. S2C,D). These data argue that the progressive hearing phenotype in the *jordan* mouse was due to postnatal stereocilia bundle degradation. Overall, we conclude that the *jordan* allele causes hearing loss by interfering with both initial elongation of stereocilia, and their postnatal maintenance. This was consistent with the p.D1647G mutation targeting the motor domain of both MYO15 isoforms (Fang et al., 2015).

### MYO15 traffics the elongation complex in *jordan* hair cells

Hair bundle development requires MYO15 dependent trafficking of EPS8, WHRN, GNAI3 and GPSM2 (the ‘elongation complex’), and mutations that prevent MYO15 trafficking (*i.e. shaker-2*) cause a short hair bundle phenotype (Belyantseva et al., 2005; Manor et al., 2011; Mauriac et al., 2017; Tadenev et al., 2019). Our finding that stereocilia lengths in the *jordan* mouse were only marginally longer than the *shaker-2*, led us to hypothesize that MYO15 trafficking was defective in *jordan* hair cells. We used the previously validated pan-MYO15 antibody PB48 (Fig. 1A), that binds an epitope common to all isoforms (Liang et al., 1999; Fang et al., 2015), to detect MYO15 in fixed cochleae from *Myo15^jd/jd^* mutants and *Myo15^+/jd^* littermates at P14. As expected, in *Myo15^+/jd^* mice, PB48 labelling was concentrated at the tips of all stereocilia rows in IHCs (Fig. 3A) (Belyantseva et al., 2003, 2005; Rzadzinska et al., 2004). We confirmed that PB48 did not label the short stereocilia of *Myo15^sh2/sh2^* IHCs at P14 (Fig. 3A), consistent with MYO15 being absent from the bundle (Belyantseva et al., 2003). In stark contrast with the *shaker-2*, we observed PB48 labelling at the tips of IHC stereocilia in *Myo15^jd/jd^* mice at both P7 and P14 (Fig. 3A,B). These data support the *jordan* and *shaker-2* mutations affecting stereocilia growth by profoundly different mechanisms: the *shaker-2* mutation blocks MYO15 trafficking, whilst the *jordan* mutation allows MYO15 to traffic but prevents stereocilia elongation.

**FIGURE 3.**
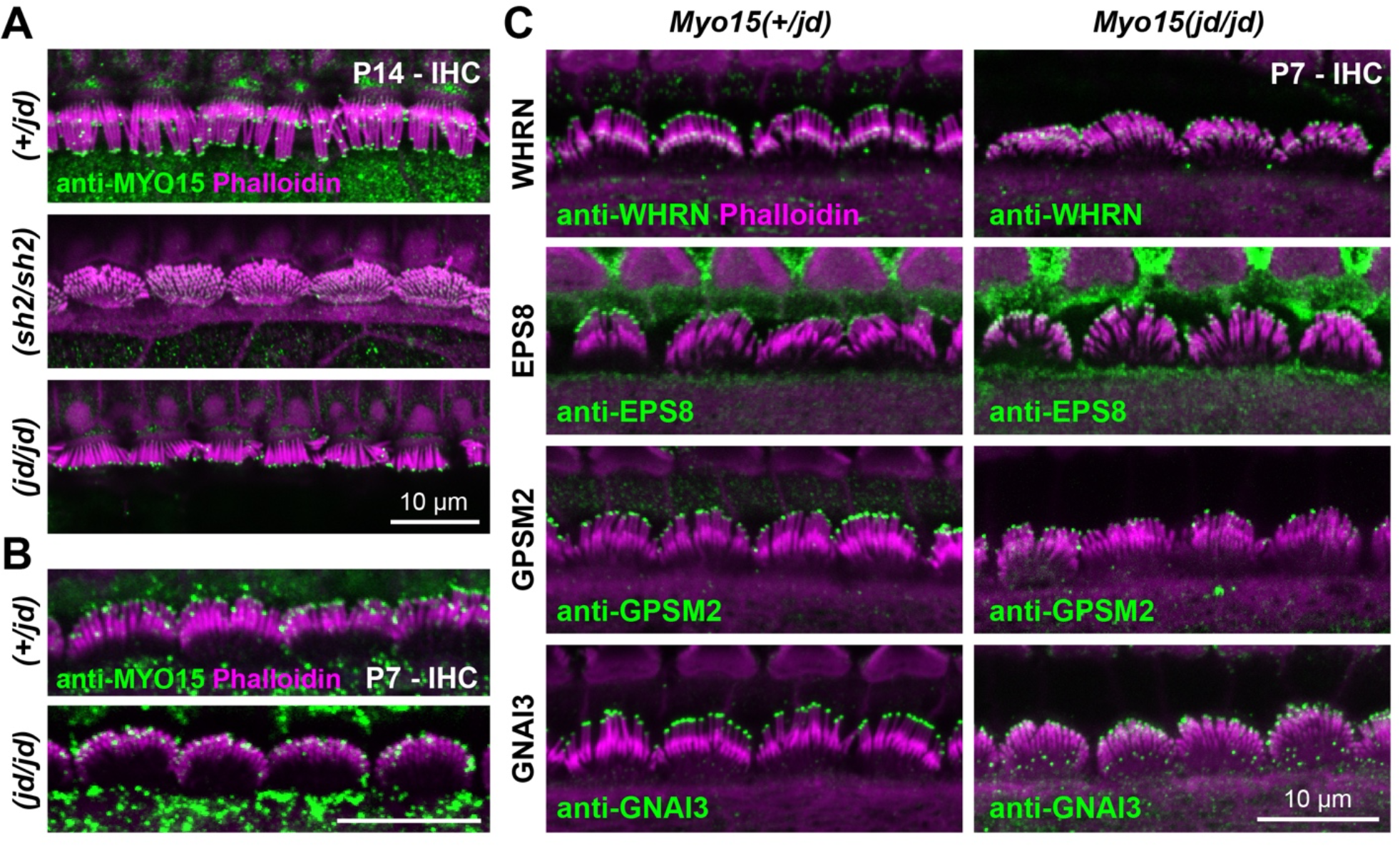
**A,B)** Immunofluorescence (IF) confocal images showing anti-MYO15 (green, PB48) in control *Myo15^+/jd^*, *Myo15^sh2/sh2^* and *Myo15^jd/jd^* IHCs fixed at P14 (A), or P7 (B). Phalloidin was used to label F-actin (magenta). **C)** IF confocal images of elongation complex proteins (green) in control *Myo15^+/jd^* and *Myo15^jd/jd^* IHCs fixed at P7, overlaid with phalloidin labelled F-actin (magenta). Images are representative of data from at least two independent animals per genotype / antibody, and are shown as maximum intensity projections reconstructed from z-stacks. Scale bars, 10 µm.

We next investigated if the *jordan* mutation interfered with trafficking of the elongation complex proteins. To test this, mutant *Myo15 ^jd/jd^* and control *Myo15^+/jd^* littermate cochleae at P7 were labelled with validated antibodies to localize WHRN, EPS8, GPSM2 and GNAI3. All of the elongation complex proteins were concentrated at the tips of the tallest stereocilia (row 1) in control *Myo15^+/jd^* hair cells (Fig. 3C), in agreement with previous work (Mburu et al., 2003; Manor et al., 2011; Zampini et al., 2011; Tarchini et al., 2016; Mauriac et al., 2017). All four proteins in the elongation complex were also targeted to the stereocilia tips in mutant *Myo15^jd/jd^* hair cells (Fig. 3C), although the signal-to-noise of GPSM2 and GNAI3 was lower, indicating a reduced overall accumulation (Fig. 3C). In older animals at P14, EPS8 and WHRN were still correctly targeted in *Myo15^jd/jd^* hair cells, however GNAI3 and GPSM2 were no longer concentrated at stereocilia tips (Fig. S3). These data are consistent with GPSM2-GNAI3 operating as a distinct complex with MYO15-EPS8 (Tadenev et al., 2019). We conclude that the initial defective stereocilia elongation in *jordan* hair cells evident at P7 was not due to gross disruption of elongation complex trafficking by MYO15.

### Actin barbed end capping is unaffected in *jordan* mutant hair cells

The presence of the elongation complex at the stereocilia tips of *jordan* hair cells suggested that the proteins known to stimulate growth were present, but insufficient to drive elongation. We considered whether an overall inhibition of actin polymerization was preventing stereocilia elongation in *jordan* hair cells. Capping proteins (CAPZ, TWF2) are present within stereocilia and regulate filament polymerization by blocking barbed end elongation (Peng et al., 2009; Avenarius et al., 2017). To detect free barbed ends, we monitored the incorporation of TMR-labelled actin monomers in permeabilized hair cells at P6. In control *Myo15^+/jd^* hair cells, TMR-actin was concentrated at row 2 stereocilia tips revealing uncapped barbed ends (Fig. 4A). TMR-actin was not consistently detected at row 1 tips, arguing that barbed ends were inaccessible at that location (Fig. 4A). The *jordan* mutation did not alter this distribution of free barbed ends and we observed a similar incorporation of TMR-actin in *Myo15^jd/jd^* hair cells compared to controls (Fig. 4A). In striking contrast, TMR-actin was incorporated at the tips of all stereocilia rows in *Myo15^sh2/sh2^* hair cells, indicating the abnormal presence of barbed ends in row 1, and an overall loss of hair bundle row identity (Fig. 4A). In summary, *jordan* hair cells retain row identity at P6, consistent with MYO15 correctly targeting the elongation complex to the stereocilia tip compartment. These data argue that inappropriate actin filament capping is unlikely the cause of short stereocilia in *jordan* hair cells. Instead, we hypothesized that a stimulatory factor independent of the elongation complex was missing.

**FIGURE 4.**
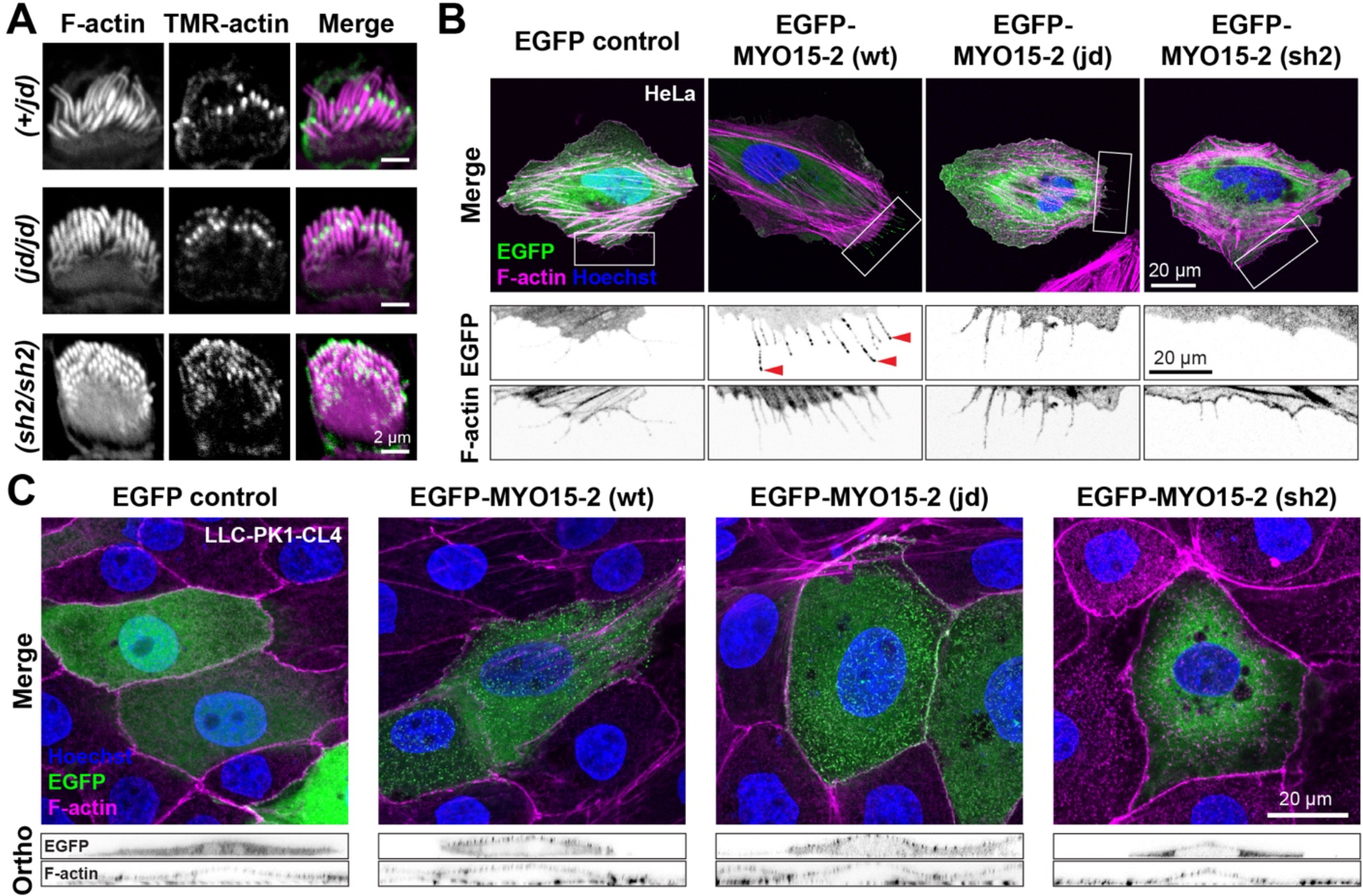
**A)** Actin barbed-end assay in detergent-permeabilized hair cells from mouse cochlear explant cultures at P6. TMR-labelled G-actin (green) was added prior to fixation to identify uncapped barbed ends. Phalloidin labelling of F-actin (magenta) is overlaid. In both *Myo15^jd/jd^* and littermate *Myo15^+/jd^* controls, barbed-ends were detected at row 2 stereocilia tips, and at the tips of all stereocilia rows in *Myo15^sh2/sh2^* hair cells. **B)** HeLa cells were transfected with EGFP-tagged *Myo15-2* expression constructs or EGFP alone (green) as indicated, fixed and probed with phalloidin (magenta) and Hoechst (blue). EGFP-M15 (wt) trafficked to filopodia tips (red arrowheads), while EGFP-M15 (jd) and EGFP-M15 (sh2) did not. Boxed regions are magnified (inverted grayscale). **C)** CL4 cells were transfected with EGFP-tagged *Myo15-2* and prepared as above. Both EGFP-M15 (wt) and EGFP-M15 (jd) concentrated at microvillar tips, while EGFP-M15 (sh2) did not. Orthogonal projections are shown (inverted grayscale). Images are representative from at least three independent experiments. Scale bars, 2 µm (A); 20 µm (B+C).

### The *jordan* mutation affects the interaction of MYO15 with actin filaments

The *jordan* missense substitution is in the MYO15 motor domain helix-loop-helix (HLH) motif that forms part of the direct binding interface with the actin filament (Gong et al., 2021). We hypothesized that a defect in MYO15’s interaction with actin might underlie the *jordan* phenotype. To explore this, we examined MYO15-2 trafficking along filopodia; actin-based structures that protrude from the periphery of heterologous cells (Belyantseva et al., 2003, 2005). We focused on MYO15-2 as the isoform responsible for stereocilia growth (Fang et al., 2015). In transfected HeLa cells, EGFP-tagged wild-type MYO15-2 accumulated at filopodia tips indicating robust anterograde myosin movement along the filopodia shaft (Fig. 4B). Discrete puncta of MYO15-2 were observed along the filopodia shaft, arising from retrograde actin filament treadmilling (Belyantseva et al., 2005; Kerber et al., 2009; Bird et al., 2017). EGFP alone did not accumulate within filopodia, proving this distribution required active myosin motility (Fig. 4B). In cells expressing the MYO15-2 *jordan* mutant, EGFP was observed diffusely along filopodia shafts and was not concentrated at filopodia tips (Fig. 4B). This was qualitatively similar to cells expressing the MYO15-2 *shaker-2* mutant (Fig. 4B), which was previously shown to not traffic along filopodia (Belyantseva et al., 2005). The inability of the MYO15-2 *jordan* mutant to traffic within filopodia and accumulate at filopodia tips was unexpected and contrary to MYO15 protein accumulating at the stereocilia tips of *Myo15^jd/jd^* hair cells (Fig. 3A).

Myosin motors are sensitive to actin filament topology (Nagy et al., 2008; Brawley and Rock, 2009; Reymann et al., 2012; Ropars et al., 2016) and we hypothesized that filopodia might not contain the appropriate repertoire of actin-binding proteins (ABPs) to support motility. To test this hypothesis, we used the porcine LLC-PK1-CL4 (CL4) epithelial cell line that generates microvilli and is a more accurate model for stereocilia (Zheng et al., 2010). In CL4 cells transfected with wild-type EGFP-MYO15-2, EGFP positive puncta localized at the tips of microvilli (Fig. 4C, orthogonal projections). In striking contrast to HeLa cells, the MYO15-2 *jordan* mutant also concentrated into microvilli and was indistinguishable from the wild-type in CL4 cells (Fig. 4C). No microvillar accumulation of the MYO15-2 *shaker-2* mutant was observed (Fig. 4C). We conclude that whilst *jordan* and *shaker-2* mutants are both immobile within filopodia, the *jordan* mutant can still actively concentrate in microvilli. These data mimic our findings from hair cells *in vivo*, and support a change in actin binding as being central to the stereocilia growth defect phenotype observed in *jordan* hair cells.

### The ATPase activity of MYO15 is altered by the *jordan* mutation

To study the interaction between MYO15 and actin filaments in more detail, we characterized the influence of the *jordan* mutation upon ATPase mechanochemistry (Bird et al., 2014; Jiang et al., 2021). Motor domain proteins were expressed in *S. frugiperda* (*Sf*9) insect cells and purified by chromatography (Fig. 5A,C). In contrast to the *shaker-2* variant, which aggregated within *Sf*9 cells, both wild-type and *jordan* motor domains were soluble. Using size exclusion chromatography (SEC), we found that FLAG-purified *shaker-2* motor domain eluted close to the void volume (Fig. 5B). In contrast, the *jordan* motor domain eluted at a similar delay volume to the wild-type (Fig. 5B), which is monomeric (Bird et al., 2014). We conclude that the *jordan* variant does not affect folding, whilst the *shaker-2* caused misfolding, consistent with the mutation being within the ATPase transducer core (Gong et al., 2021).

**FIGURE 5.**
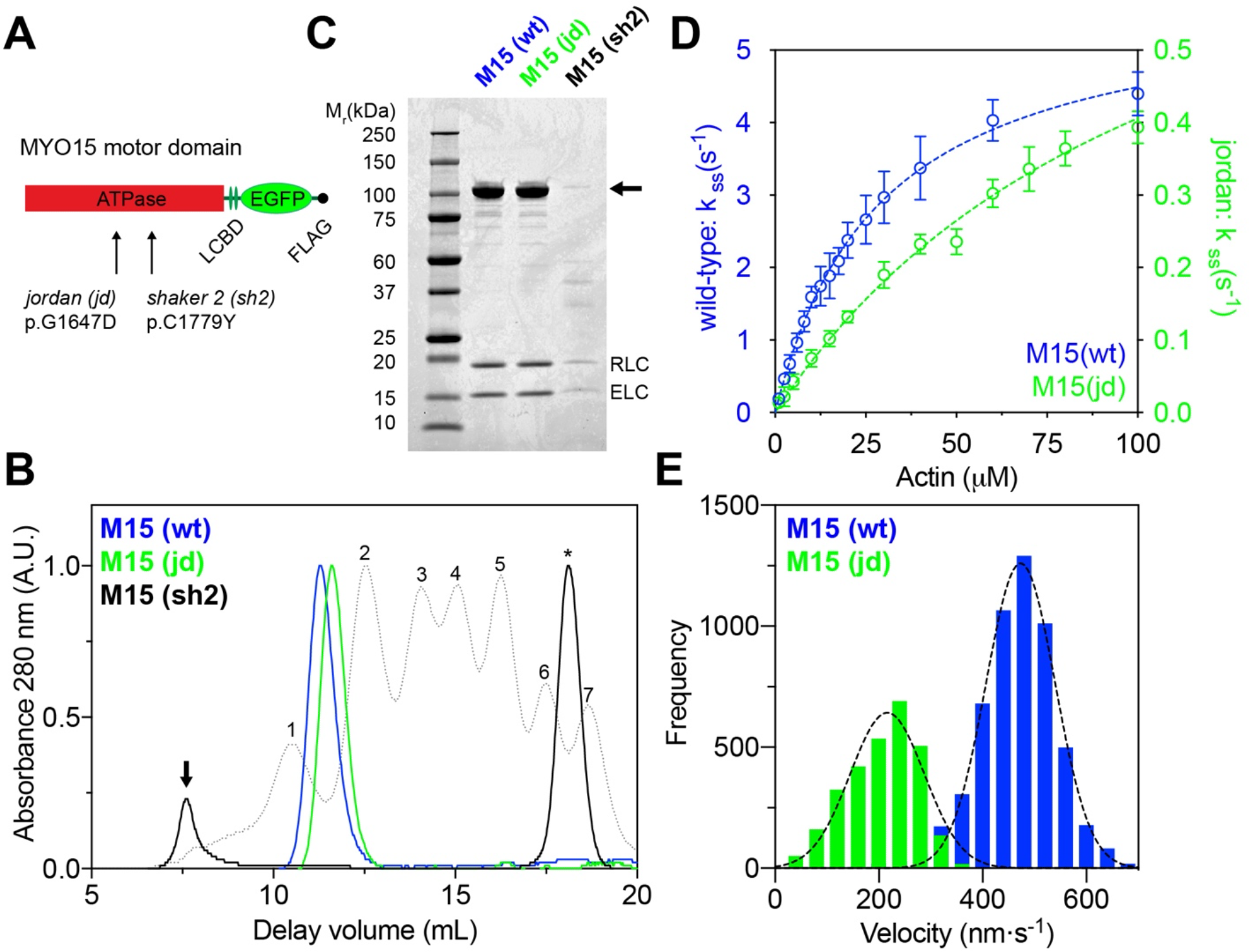
**A)** Cartoon of MYO15 motor domains expressed in *Sf*9 cells, consisting of the ATPase and two light chain binding domains (LCBD). **B)** Size exclusion chromatography analysis of FLAG/IEX purified M15(wt) and M15(jd) proteins. M15(sh2) elutes close to the void volume (arrow) and was aggregated. Protein calibration standards (dotted lines), FLAG peptide (asterisk). **C)** SDS-PAGE analysis of SEC purified motor domain proteins. The motor domain (arrow) co-purifies with RLC and ELC light chains for all variants. M15(sh2) was misfolded and extracted from *Sf*9 cells at low yield. **D)** Steady-state ATPase activation of M15(wt) and M15(jd) motor domains measured using a NADH-coupled assay at 20 ± 0.1°C. Reactions were performed with [F-actin] as shown. Rectangular hyperbola fits are shown for M15(wt) (blue, *k*_cat_ = 5.8 ± 0.2 s^−1^, *k*_ATPase_ = 29.1 ± 2.1 μM) and for M15(jd) (green, *k*_cat_ = 0.87 ± 0.04 s^−1^, *k*_ATPase_ = 114.3 ± 8.2 μM, n = 4). **E)** Frequency histogram of F-actin velocities in a gliding filament assay at 30 ± 0.1°C. Gaussian fits (dotted line) are overlaid for M15(wt) (473 ± 67 nm⋅s^−1^, n = 5449 filaments, mean ± SD) and M15(jd) (216 ± 71 nm⋅s^−1^, n = 2844 filaments). Data are from 2 independent experiments.

To test for differences in enzymatic activity between wild-type and *jordan* motor domain variants (Fig. 5A), we measured steady-state ATP hydrolysis using an enzyme-linked NADH assay. The basal ATPase activity (*k*_basal_) of the wild-type motor domain was 0.06 ± 0.01 s^-1^, measured without F-actin. The addition of F-actin caused a 97-fold increase in ATPase activity to an extrapolated maximum catalytic rate (*k*_cat_) of 5.8 ± 0.2 s^-1^ (Fig. 5D). Half-maximal activation of the ATPase activity (*K*_ATPase_) was reached at 29.1 ± 2.1 µM F-actin, as previously reported (Bird et al., 2014; Jiang et al., 2021). Using identical assay conditions, the basal ATPase rate of the *jordan* variant was unchanged at *k*_basal_ = 0.07 ± 0.01 s^-1^. Strikingly, there was a reduced 7-fold maximal activation to *k*_cat_ = 0.87 ± 0.04 s^-1^ (Fig. 5D). Furthermore, half-maximal ATPase activation was increased to *K*_ATPase_ = 114.3 ± 8.2 µM actin, signifying a 4-fold reduction in the apparent actin affinity compared to wild-type.

The mechanical activity of wild-type and *jordan* motor domains was measured using an *in vitro* gliding filament assay, where actin filaments are propelled across a microscope cover glass functionalized with motor domain protein (Sellers et al., 1993). Due to the reduced apparent affinity of the *jordan* motor domain for actin, we lowered the salt concentration to 10 mM KCl in these assays. The wild-type motor domain robustly propelled actin filaments at 473 ± 67 nm·s^-1^ (Fig. 5E), consistent with previous data (Bird et al., 2014). In contrast, the *jordan* mutant moved filaments at 216 ± 71 nm·s^-1^, a 2-fold reduction from the wild-type velocity (Fig. 5E). Overall, we conclude that whilst the *jordan* mutation caused a significant ATPase defect and decreased the motor domain’s apparent affinity for actin, the motor domain was still mechanically active. Our data further establish a functional correlation between motor domain activity and the severity of hearing loss in a *Myo15* allelic series. The *jordan* motor domain retained partial activity consistent with this mutation causing intermediate hearing loss, whilst the *shaker-2* motor domain was misfolded and associated with the most severe phenotype.

### The MYO15 motor domain directly stimulates actin polymerization

Our *in vitro* analysis of purified MYO15 motor activities were consistent with the *jordan* mutant maintaining the ability to traffic the elongation complex and accumulate at stereocilia tips. As stereocilia nevertheless fail to grow normally in *jordan* hair cells, we hypothesized that independent of molecular trafficking, MYO15 had an additional function at the stereocilia tip critical for elongation. Classic biochemical studies of muscle myosin have shown its motor domain (*e.g.* subfragment-1, S1) can directly stimulate actin polymerization *in vitro*; however, the physiological relevance of this activity *in vivo*, if any, is unknown (Yagi et al., 1965; Miller et al., 1988; Fievez and Carlier, 1993; Lheureux et al., 1993). We hypothesized that the MYO15 motor domain could directly stimulate actin polymerization to drive stereocilia elongation, and that this critical activity is disrupted by the *jordan* deafness mutation.

We tested this hypothesis using pyrene-conjugated globular actin (G-actin) monomers that increase in fluorescence as they polymerize into filamentous actin (F-actin) (Cooper et al., 1983). As a control, 2 µM G-actin was polymerized with 1x KMEI (50 mM KCl, 1 mM MgCl2, 1 mM EGTA, 10 mM imidazole) while monitoring pyrene fluorescence. F-actin assembled with an initial lag phase representing the kinetically unfavourable nucleation step (Fig. 6A, red line). When the reaction was repeated with 1 µM wild-type motor domain + 2 µM G-actin + KMEI, actin polymerization initially followed the same trajectory (Fig. 6A, blue line). However, following a delay of ∼ 400 seconds, an inflection was observed where the rate of actin polymerization reduced momentarily (Fig. 6A, blue line), before rapidly increasing and reaching steady-state (Fig. 6A,B). We tested if MYO15-stimulated actin polymerization was sensitive to the *jordan* mutation. Under identical experimental conditions using 1 µM *jordan* MYO15 motor domain, the actin polymerization rate was initially indistinguishable from the control reaction. However, following an inflection at ∼ 600 seconds, the actin polymerization rate decreased below that of the G-actin alone control (Fig. 6A). We conclude that the wild-type MYO15 motor domain stimulated actin polymerization, whilst the deafness-causing *jordan* mutant blocked this activity.

**FIGURE 6.**
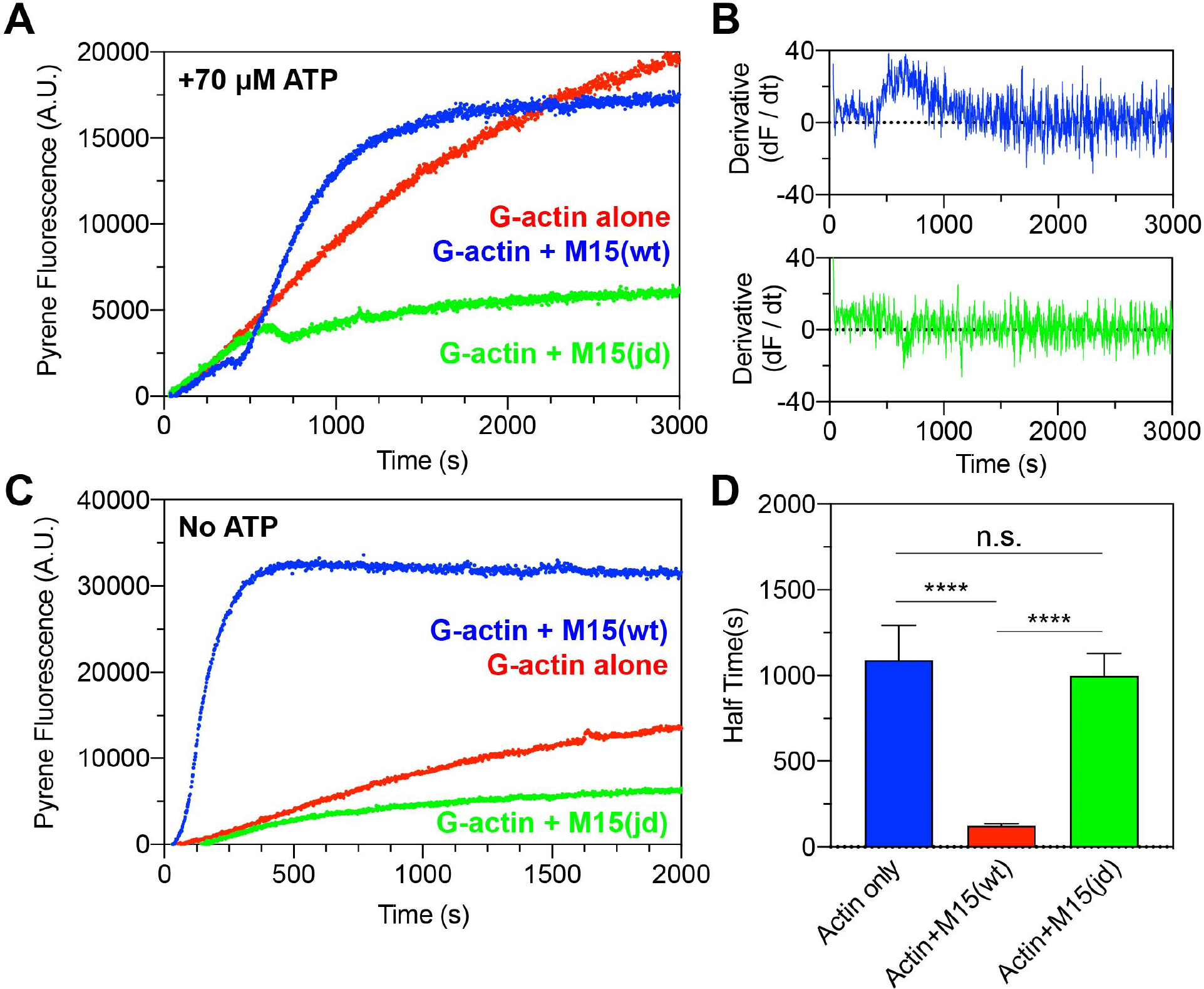
**A)** Fluorescent time-course of 2 µM G-actin (10% pyrene) measured in a fluorimeter (red trace). Polymerization was induced by addition of KMEI (1x) at t = 0 s. Introduction of 1 µM M15(wt) (blue) at t = 0 s increases in actin polymerization following an inflexion point (arrow), whilst 1 µM M15(jd) (green) does not (arrow). Reactions contains 70 µM ATP carried over from the actin storage G-buffer. **B)** First derivative of pyrene fluorescence with respect to time for G-actin + M15(wt)(blue) and G-actin + M15(jd)(green). **C)** Fluorescent time course of 2 µM G-actin (10% pyrene) induced to polymerize by KMEI buffer in the absence of free ATP (red trace). G-actin monomers were desalted immediately prior to use, ensuring they were ATP bound. Addition of 1 µM M15(wt) (blue) triggers an almost immediate increase in actin polymerization. **D)** Quantification of time to reach half maximal fluorescence (n = 4).

With either wild-type or *jordan* motor domains, we observed a delay before the actin polymerization rate deviated from the control, with the delay longer for *jordan* (∼ 600s) versus wild-type (∼ 400s). We hypothesized this difference arose from residual ATP (70 µM) being introduced from the G-actin storage buffer; ATP binding would prevent the motor domain from occupying the strongly actin bound (rigor) state. Furthermore, we expected this residual ATP to be hydrolysed faster by the wild-type motor domain, compared with the *jordan* mutant (Fig. 5D). To test this, we repeated the pyrene polymerization assay in the absence of free ATP. To ensure G-actin monomers were ATP-bound, G-actin was desalted immediately prior to use to remove free ATP from solution. Using this approach, we measured the polymerization of 2 µM G-actin + KMEI in the absence of free ATP (Fig. 6C) and observed kinetics comparable to before (Fig. 6A). Strikingly, pyrene fluorescence immediately increased upon addition of 1 µM wild-type motor to the reaction, and rapidly saturated (Fig. 6C) with a significantly reduced half-time of 124 ± 12 s, compared with 1091 ± 202 s for G-actin + KMEI alone (Fig. 6D). While addition of 1 µM *jordan* motor domain to 2 µM G-actin + KMEI reduced the overall extent of actin polymerization (Fig. 6C), the half-time to saturation was 1000 ± 65 s and not significantly different to G-actin + KMEI alone (Fig. 6D). We conclude that the wild-type MYO15 motor domain strongly accelerated actin filament polymerization in the absence of ATP, implicating the rigor state as critical for this activity. Combined with experiments from our companion study showing that ADP-bound MYO15 also stimulated polymerization (Gong et al., 2021), these data identify the strong actin-binding states as the key MYO15 ATPase intermediates that stimulate actin polymerization.

### The *jordan* deafness mutation inhibits *de novo* nucleation of actin filaments

To directly visualize how the MYO15 motor domain accelerates polymerization, we performed actin polymerization assays using total internal reflection fluorescence microscopy (TIRFM) where the elongation of individual filaments can be tracked (Fujiwara et al., 2002). A control time-lapse of 1 µM G-actin + KMEI revealed a slow rate of filament nucleation, with short polymers attaching to surface and elongating (Fig. 7A, top row). We repeated the experiment with 1 µM wild-type motor domain and observed a large burst in filament density (Fig. 7A, middle row) that was statistically significant at 480 seconds when compared to the actin-alone control (Fig. 7B). In contrast, when polymerization experiments were repeated with 1 µM *jordan* motor domain (Fig. 7A, bottom row), there was no increase in actin filament density above the actin-alone control (Fig. 7C). Using kymographs to track filament barbed ends, we found that the motor domain could influence actin filament elongation rates. In control experiments, barbed ends elongated at 16.2 ± 2.6 nm·s^-1^ (Fig. 7C,D), consistent with previous data using TMR-conjugated G-actin on Cys374 (Kuhn and Pollard, 2005). Elongation rates were significantly reduced by the addition of either wild-type or *jordan* motor domain, to 13.3 ± 4.3 nm·s^-1^ and 13.4 ± 3.5 nm·s^-1^, respectively (Fig. 7C,D). The decrease in elongation rate was dependent upon timing of the filament burst (Fig. 7B). When data were binned as pre-burst (< 320 s), the addition of wild-type or *jordan* motor domains did not significantly alter elongation rates compared to the control (Fig. 7E). In contrast, the presence of either wild-type or *jordan* motor domain post-burst (> 720s) both significantly, and equally, reduced elongation rates below the actin alone control (Fig. 7F).

**FIGURE 7.**
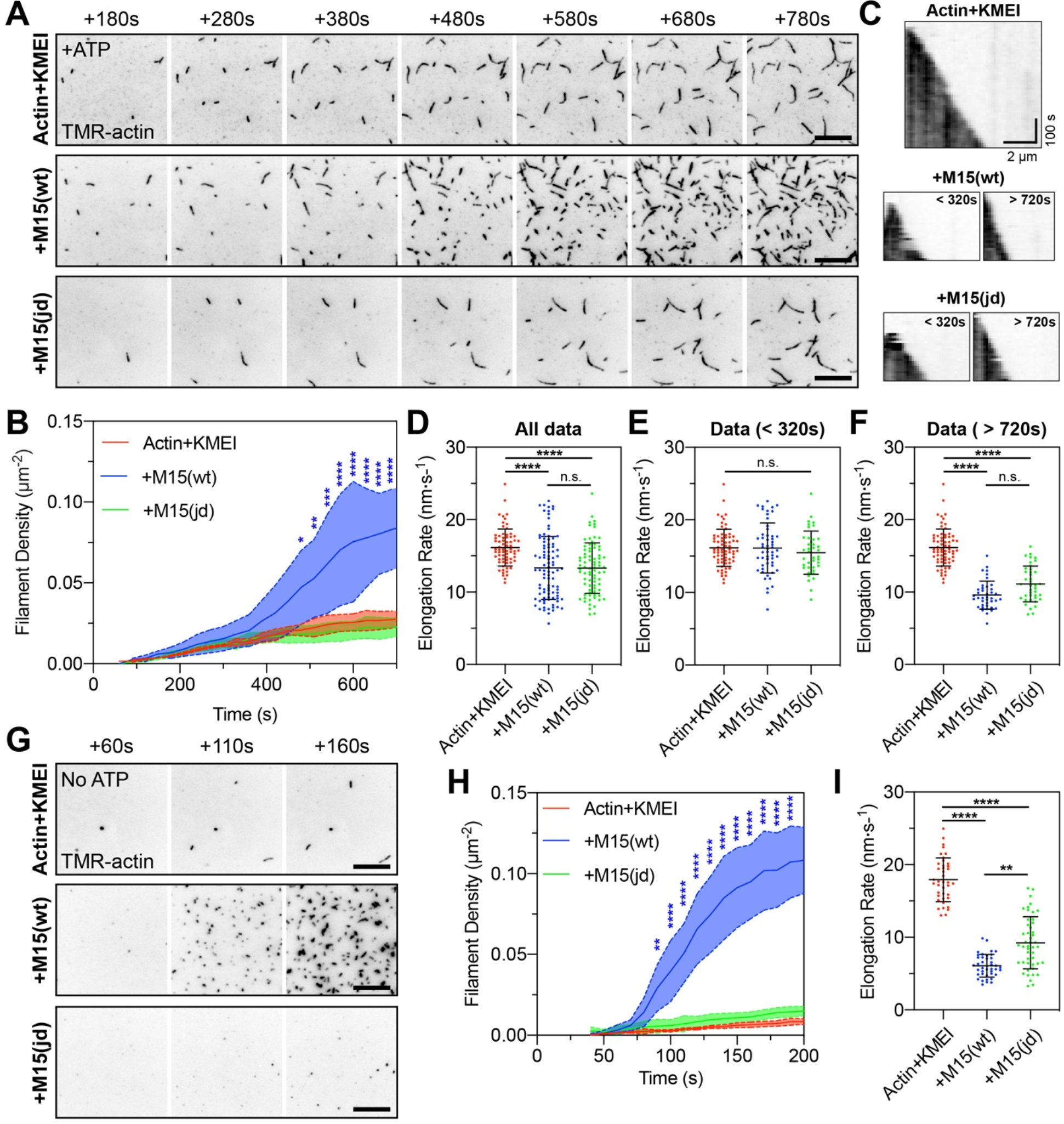
**A)** TIRFM visualization of actin filaments polymerizing on PEG-biotin-neutravidin functionalized cover glass. Polymerization of 1 µM G-actin (20% TMR + 10% biotin labelled) was induced by 1 x KMEI (50 mM KCl, 1 mM MgCl2, 1 mM EGTA, 10 mM imidazole, pH 7.0) at t = 0 s, in the presence of 25 µM ATP. Representative time-lapses shown for: 1 µM G-actin (top), 1 µM G-actin + 1 µM M15(wt) (middle), and 1 µM G-actin + 1 µM mutant M15(jd) (bottom). **B)** Quantification of actin filament density shows delayed nucleation activity of MYO15 in the presence of ATP. **C)** Kymographs of actin filament elongation. **D**) Barbed end elongation rates for G-actin alone (red), G-actin + M15(wt) (blue), G-actin + M15(jd) (green). **E + F)** Elongation rate data (from D) re-binned as before (E), or after (F) nucleation. The G-actin + KMEI control data set (from D) is reproduced identically as a comparator in E and F. **G)** Time-lapse of actin filaments polymerization induced by KMEI at t = 0 s, with no ATP in solution. G-actin (ATP) monomers were prepared by desalting into ATP-free G-buffer. **H)** Actin filament density shows nucleation activity of MYO15 is accelerated in the absence of ATP. **I)** Barbed end filament rates in the absence of free ATP. Reaction deadtimes were typically 50 s and included in quantification. TIRFM images are shown as inverted grayscale. Scale bars are 10 µm (A,G). Data are from 4 experiments, using 2 independent protein preparations.

Nucleation by wild-type MYO15 took an average of 480 seconds to be significantly increased above the spontaneous nucleation rate of G-actin + KMEI alone (Fig. 7B). We considered whether the delayed production of short filaments was caused by MYO15 severing actin filaments that nucleated spontaneously. To test this hypothesis, we repeated our experiments in the absence of free ATP to force the motor domain into rigor binding and accelerate polymerization. Free ATP was desalted from 1 µM G-actin monomers; this did not affect barbed-end elongated rates when polymerized by KMEI (Fig. 7D,I). As expected, when 1 µM wild-type motor domain was included in the reaction, there was a potent nucleation of short actin filaments observed within 90 s (Fig. 7G,H), confirming acceleration of nucleation when the motor domain was forced into rigor. Under identical ATP free conditions, 1 µM *jordan* motor domain did not nucleate additional filaments compared with the G-actin + KMEI control (Fig. 7G,H). Similar to experiments where ATP was present, but presumably hydrolyzed (Fig. 7F, > 720 s), we saw that both wild-type and *jordan* motor domains significantly reduced barbed-end elongation rates (Fig. 7I). We conclude that the MYO15 motor domain exerts multiple direct effects on actin polymerization, including catalyzing *de novo* filament nucleation and reducing barbed end elongation rates in a nucleotide-dependent fashion. Our data show that MYO15-induced actin nucleation is a key process targeted by the *jordan* deafness mutation, and argue that this activity is critical for stereocilia elongation and hearing.

## DISCUSSION

Plasticity of the stereocilia actin core is central to hair bundle development, mechano-sensitivity and hearing. Existing evidence that MYO15 moves along stereocilia actin filaments has led to a cargo trafficking model, where the elongation complex is delivered to the stereocilia tips and provides the actin regulation machinery necessary for elongation. Here, we uncover a previously unknown activity of MYO15 that requires the cargo-trafficking model to be re-evaluated. *In vitro*, we show that MYO15 can directly stimulate actin filament nucleation and that this activity is inhibited by the *jordan* mutation, providing key evidence this activity is functionally linked to deafness pathology. *In vivo*, the *Myo15 jordan* mutation stunted stereocilia growth, leading to an early-onset, progressive hearing loss in mice. The effect of the *jordan* mutation to inhibit MYO15-induced actin nucleation *in vitro*, and block stereocilia elongation *in vivo*, provides the functional link between these two processes. We propose that in addition to trafficking key molecular cargos, MYO15 directly catalyses actin filament nucleation at the stereocilia tip, and that both these processes combined are required for normal stereocilia growth.

Using purified proteins and reconstituted assays, the MYO15 motor domain exerted multiple effects upon actin filament polymerization *in vitro*. The motor domain: 1) nucleated actin filaments *de novo*, and 2) slowed the elongation rate of existing actin filaments. No additional proteins were required, demonstrating that these properties were intrinsic to the purified actomyosin system. The *jordan* mutation completely blocked MYO15-induced actin filament nucleation, whilst filament elongation rates were unaffected relative to wild-type MYO15. We conclude that MYO15’s nucleation activity is the key polymerization effect disrupted by the *jordan* deafness mutation. Similar to MYO15, muscle myosin was shown in classic biochemical experiments to stimulate actin filament nucleation *in vitro* (Yagi et al., 1965; Miller et al., 1988; Fievez and Carlier, 1993; Lheureux et al., 1993). The MYO15 motor domain shares a common structural fold with muscle myosin (Gong et al., 2021), and although the mechanism of muscle nucleation remains controversial (Lheureux et al., 1993), we believe by homology to muscle that MYO15 nucleates actin through stabilizing inter-subunit contacts that establish the protofilament (Fievez et al., 1997a, 1997b). In our companion study (Gong et al., 2021), the MYO15 motor domain is further shown to influence structural plasticity within the DNaseI-binding loop (D-loop), a sub-domain of actin that regulates the monomer to polymer transition (Dominguez and Holmes, 2011). Combined, our data argue that nucleation is a shared function for at least some members of the myosin superfamily, and that myosin motors can directly influence cytoskeletal plasticity in addition to their canonical role generating force.

A cellular function for myosin-driven actin nucleation has, to the best of our knowledge, not previously been reported in any tissue. Our study provides evidence that MYO15-driven nucleation contributes to hair cell function *in vivo*, and furthermore is a cause of disease when disrupted. Actin nucleation by MYO15 appears distinct from other nucleation factors, such as formins, Arp2/3, Spire and Cobl (Machesky et al., 1994; Mullins et al., 1997, 1998; Sagot et al., 2002; Quinlan et al., 2005; Ahuja et al., 2007). MYO15-driven nucleation was nucleotide-sensitive, and strongly promoted nucleation under rigor (nucleotide-free) conditions, or when bound to ADP (Gong et al., 2021). Both conditions populate the motor domain into strong actin binding states (Jiang et al., 2021). Conversely, the presence of ATP in assays prevented nucleation and we hypothesize this is due to the motor domain populating a weakly actin-bound state (Jiang et al., 2021). The nucleotide-sensing properties of MYO15 may therefore couple local concentrations of ATP and ADP to actin polymerization. We speculate this activity could form the basis of a tuning mechanism, where dynamic changes in nucleotide concentrations control the height of developing stereocilia. The concentrations of ATP and ADP in stereocilia are unknown, and whilst a creatine kinase ATP generation system has been described in avian hair cells (Shin et al., 2007), its activity in mammals is unclear (Krey and Barr-Gillespie, 2019). In addition to nucleotide state, MYO15-driven nucleation *in vitro* was observed at high protein concentrations equimolar with actin, in agreement with data from muscle myosin (Miller et al., 1988). The concentration of MYO15 in hair cells is unknown, however micromolar quantities may be present at the stereocilia tips, especially as MYO15 actively traffics and self concentrates into this subcellular compartment (Belyantseva et al., 2003). Other unconventional myosin motors critical for hearing, including MYO1C, MYO3A, MYO6 and MYO7A (Avraham et al., 1995; Gibson et al., 1995; Gillespie and Corey, 1997; Hasson et al., 1997; Holt et al., 2002; Salles et al., 2009; Ebrahim et al., 2016), also concentrate in specific stereocilia sub-domains and it is exciting to consider whether they also directly regulate actin polymerization as part of their cellular function.

In parallel with identifying MYO15 as an actin nucleator, we used the *jordan* mutant mouse to probe sufficiency of the elongation complex to drive stereocilia elongation. Each protein in the elongation complex is critical for growth, but existing models do not explain their molecular activty, except that they must be delivered to the stereocilia tip via MYO15-driven trafficking. This model is based on the *shaker-2* mouse, where stereocilia growth is inhibited and both MYO15 and the elongation complex are absent from the hair bundle (Belyantseva et al., 2005; Delprat et al., 2005; Manor et al., 2011; Zampini et al., 2011; Mauriac et al., 2017; Tadenev et al., 2019). Our data show that MYO15-driven actin nucleation and elongation complex trafficking activities are simultaneously lost in the *shaker-2*, leaving their relative contributions unknown. By contrast, MYO15-driven trafficking of the elongation complex is initially (largely) preserved in the short stereocilia of the *jordan* mouse. These data argue that the elongation complex proteins are necessary, but not sufficient for driving stereocilia growth, and that MYO15-nucleation activity is critical *in vivo*. We cannot exclude the possibility of reduced MYO15 trafficking in *jordan* hair cells; indeed, gliding filament velocities of the mutant motor were ∼ 50% of the wild-type *in vitro*. However, there is no evidence that the concentration of MYO15 at the stereocilia tips determines stereocilia length. Neither the elimination of MYO15-1 from shorter stereocilia rows (Fang et al., 2015), nor the over-expression of MYO15-2 alters initial stereocilia lengths (Belyantseva et al., 2003, 2005). We also considered whether the *jordan* mutation might interfere with MYO15 binding to another protein critical for stereocilia growth. Our companion study (Gong et al., 2021) localizes the *jordan* missense substitution to the actomyosin interface, and shows a direct structural effect within the actin monomer itself. Combined with our study, these findings strongly suggest that MYO15 directly accelerates actin nucleation at the stereocilia tip by structurally regulating the actin molecule.

Our study raises new questions concerning the stability and plasticity of the stereocilia cytoskeleton, processes that must be tightly controlled for mechano-transduction and lifelong hearing (Vélez-Ortega and Frolenkov, 2019). Stereocilia actin filaments are unidirectionally polarized with their fast-growing barbed ends orientated towards the tip compartment, where new actin monomers are incorporated (Flock and Cheung, 1977; Tilney et al., 1980; Schneider et al., 2002; Drummond et al., 2015). Although actin capping proteins also contribute to stereocilia architecture (Peng et al., 2009; Avenarius et al., 2017), we find that excessive barbed end capping is not the cause of short stereocilia in *Myo15* mutant mice. Given the presence of barbed ends, why might actin nucleation be required to extend pre-existing filaments at stereocilia tips? One possible explanation is that stereocilia elongation occurs through direct end-to-end annealing of short actin polymers nucleated by MYO15, rather than through addition of individual actin monomers at the barbed end. Pre-formed actin filaments can anneal into longer filaments *in vitro* (Murphy et al., 1988; Kinosian et al., 1993; Sept et al., 1999), and more recent *in vitro* studies have detected filament extension through incorporation of short polymers, in addition to the dominant mode of monomer addition (Young et al., 2018). Interestingly, endocytosis in yeast is proposed to use end-to-end actin filament annealing rather than monomer polymerization (Okreglak and Drubin, 2010). Though speculative in hair cells, the extension of the stereocilia actin core using short polymer annealing may allow for rapid changes in stereocilia length that are likely needed to continually maintain the sensitivity of the MET machinery (Vélez-Ortega et al., 2017; Krey et al., 2020).

Given the potent ability of MYO15 to nucleate actin filament polymerization *in vitro*, we infer this activity must be tightly regulated in hair cells. There is extensive evidence for intramolecular regulation throughout the myosin superfamily (Heissler and Sellers, 2016). For example, myosin 5 (MYO5A) exists in an autoinhibited conformer where the globular tail domain binds and inhibits motor domain activity; binding of melanophilin (MLPH) to MYO5 releases this autoinhibition to activate the motor domain (Li et al., 2005; Liu et al., 2006; Thirumurugan et al., 2006; Sato et al., 2007; Sckolnick et al., 2013). Similar autoinhibitory regulation mechanisms have been described for other members of the MyTH4 - FERM myosin family, *e.g.*, MYO7A and MYO10 (Umeki et al., 2009, 2011; Yang et al., 2009; Sakai et al., 2011). We hypothesize that part of the elongation complex function may be to bind MYO15 and shift the equilibrium between active and inhibited states. Intramolecular regulation may also explain how different MYO15 isoforms independently exert their effects upon hair bundle architecture (Fang et al., 2015). MYO15-1 and MYO15-2 possess an identical motor domain competent to regulate actin polymerization, but differ by the addition of a 133 kDa N-terminal domain (Fig. 1A). How the N-terminal domain affects the actin nucleation activity of the motor domain is an important future question and will help uncover how the MYO15-1 isoform controls actin polymerization in shorter stereocilia rows with active MET (Fang et al., 2015). In conclusion, we reveal a new function for MYO15 and argue that defective nucleation of actin polymerization interferes with stereocilia architecture and causes progressive hearing loss. We speculate that defective actin nucleation by MYO15 is a central molecular pathology underlying DFNB3 hereditary hearing loss in humans.

## MATERIALS AND METHODS

### Mice

Pedigree MPC190 was identified from a phenotype-driven mutagenesis screen undertaken at the MRC Harwell Institute (Potter et al., 2016). Briefly, *N*-ethyl-*N*-nitrosourea (ENU) mutagenized C57BL/6J males were mated with wild-type ‘sighted C3H’ (C3H.Pde6b^+^) females. Resulting G1 males were crossed with C3H.Pde6b^+^ females to produce G2 females, which were screened for the *Cdh23^ahl^* allele. *Cdh23^+/+^* G2 females were backcrossed to their G1 fathers to generate recessive G3 pedigrees, which entered a longitudinal phenotyping pipeline that included click box and ABR tests to assess auditory function (Hardisty-Hughes et al., 2010). DNA from mice exhibiting hearing loss, and normal hearing pedigree mates, was prepared from ear biopsies and used for linkage mapping utilizing the Illumina GoldenGate Mouse Medium Density Linkage Panel (Gen-Probe Life Sciences Ltd, UK), which identified a critical interval on chromosome 11. DNA was extracted from mouse MPC190/2.18a and subject to whole genome sequencing employing the Illumina HiSeq platform (Oxford Genomics Centre, Wellcome Trust Centre for Human Genetics). Subsequent alignment to the reference genome (NCBIM38/mm10) identified a homozygous, non-synonymous coding lesion in the *Myo15* gene.

At the MRC Harwell Institute, mice were housed and maintained at the Mary Lyon Centre under conditions outlined in the Home Office Code of Practice, with all animal procedures licenced by the Home Office under the Animals (Scientific Procedures) Act 1986, UK and approved by the local Ethical Review Committee (PBF9BD884 to MRB). At MRC Harwell, *jordan* mice were crossed to C57BL/6N (*Cdh23^753A>G^*) ‘repaired’ mice (Mianné et al., 2016) until congenic. Concurrently, *jordan* mice were imported to the NIH and the University of Florida (UF) and maintained on a ‘sightless C3H’ (C3H.Pde6b^rd1^) background. Animal procedures were approved the Institutional Animal Care and Use Committees (IACUC) at UF (#201910739 to JEB) and at the NIH (#1263-15 to TBF). Genomic DNA from mouse tail biopsies was used as template in a PCR reaction with primers (5’-CAGGAGGAGTACATCCGGG-3’, 5’-AGACCACAGAAGTATCTGGGTCTT-3’). The resulting 161 bp amplicon was analyzed by *MlsI* endonuclease digestion. Resulting restriction fragments lengths unambiguously detected wild-type (161 bp) and mutant (116 bp + 45 bp) alleles.

### Auditory phenotyping + behavioral testing

Auditory Brainstem Response (ABR) were recorded as previously described (Dunbar et al., 2019). Briefly, mice were anaesthetised via intraperitoneal injection with ketamine hydrochloride (100 mg·kg^-1^) and xylazine (10 mg·kg^-1^). Anesthetized mice were placed inside a sound-attenuated chamber (ETS-Lindgren) and recording electrodes (Grass Telefactor F-E2-12) placed sub dermally over the vertex (active), right mastoid (reference), and left flank (ground). ABR responses were collected using TDT system III hardware and BioSig software (Tucker Davis Technology). Stimuli were presented free field from a ES1 transducer (TDT) calibrated at a distance of 1 cm from the right ear. Stimuli consisted of 0.1 ms broadband clicks or 7 ms tone-bursts at 8 kHz, 16 kHz and 32 kHz. Each stimulus was presented at a maximum 90 dB SPL, followed by decreasing steps of 5 - 10 dB SPL until no replicable response peaks were observed. ABR operators were blind to genotype.

Distortion Product Oto-Acoustic Emissions (DPOAE) were recorded as a terminal procedure in 12 weeks old mice, as described (Dunbar et al., 2019). Mice were anaesthetised with a modified ketamine/xylazine solution (see ABR), with the addition of acepromazine (2 mg·ml^-1^, 8% v/v). Anesthetized mice had a section of the pinna removed to access the external auditory meatus. Mice were placed inside a sound-attenuated chamber (ETS Lindgren), and the DPOAE probe assembly was inserted into the ear canal. In-ear calibration was performed before each test. An ER10B+ low-noise probe microphone (Etymotic Research) was used to measure DPOAEs near the tympanic membrane. Tone stimuli were presented via separate MF1 (Tucker Davis Technology) speakers, with f1 and f2 at a ratio of f2/f1 = 1.2 (L1 = 65 dB SPL, L2 = 55 dB SPL). The f1 and f2 tones were presented continuously in specific tone-bursts between 8 and 32 kHz.

### Scanning electron microscopy

Inner ears were dissected and fixed overnight at 4°C in 0.1 M phosphate buffer, 2.5% glutaraldehyde (Sigma-Aldrich). Post-fixed ears were decalcified in 4.3% EDTA in phosphate buffer at 4°C, before dissecting out the sensory epithelium. Samples were processed with alternating 1% osmium tetroxide (Agar Scientific) in 0.1 M sodium cacodylate (Sigma-Aldrich) and 1% thiocarbohydrazide (Sigma-Aldrich) treatments. Osmicated samples were dehydrated in graded ethanols (25% to 100%) at 4°C and stored in 100% acetone until critical point drying with liquid CO₂ (EM CPD300, Leica Microsystems Ltd.). Samples were mounted onto stubs using silver paint (Agar Scientific), sputter-coated with platinum (Q150R S, Quorum Technologies) and visualised with a scanning electron microscope (JSM-6010LV, JEOL).

For morphometric analyses, image pairs from the middle cochlear turn IHC and OHC bundles were captured with a 5° tilt angle difference at a constant working distance of 20 µm. Stereocilia from at least two bundles per animal were measured using ImageJ (https://imagej.nih.gov), with a minimum of 14 stereocilia for each condition. Length measurements were taken from the rear aspect of the hair bundle, so that the length of the tallest stereocilia (row 1) could be measured from the cuticular plate insertion to the tip. Estimates for actual stereocilia length were calculated using a pseudo-eucentric tilting approach (Bariani et al., 2005). A single measure x_1_ (length of stereocilium) was taken from the first image and measured again (x_2_) on the corresponding 5°-tilted image pair. Perpendicular counter-measures (y_1_ and y_2_), matched to (x_1_ and x_2_) were also recorded. Equation 1 was used to estimate uncertainty (ζ) due to plane rotation. Using the uncertainty estimate ζ from Equation 1, every pair of tilted measures (x_1_ and x_2_) was processed using Equation 2 to obtain a close approximation (ξ) of the true length of each stereocilia measured.

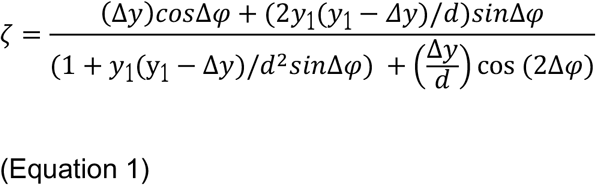

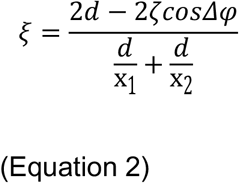

Where: ζ = uncertainty estimate, ξ = estimate of true size of structure of interest, x_1,2_ = tilted paired-measures of structure of interest, y_1,2_ = perpendicular counter-measures to measures x_1,2_, Δy = arithmetic difference of counter-measures y_1_ and y_2_, Δφ = tilting angle (5°), *d* = working distance (20µm).

In addition, low-magnification (1500 - 2000X) fields of mid-turn sensory epithelium were imaged and the OHC bundles counted and visually categorised as either: ‘Intact’, ‘Abnormal / Damaged’ where the bundle had an abnormal shape or was missing stereocilia, or ‘Missing Bundle’ where a cuticular plate was observed with no stereocilia. Cochleae from four different animals per genotype were examined, with a total of 1344 wild type and 972 *jordan* OHC bundles.

### Whole mount immunofluorescence and confocal microscopy

Mouse inner ears were fixed in 4% paraformaldehyde (EMS Diasum) in PBS for 2 hours at room temperature, washed in PBS, and then micro-dissected to isolate the cochlear sensory epithelium. Samples were permeabilized in 0.5% (v/v) Triton X-100 in PBS for 30 minutes at RT, followed by blocking in 5% normal goat serum (Sigma-Aldrich), 2% bovine serum albumin (Fraction V, Roche) in PBS for 1 hour at RT. Primary antibodies were diluted in blocking solution and incubated with samples at 4°C overnight. Primary IgG antibodies used were: PB48 rabbit anti-MYO15 (Liang et al., 1999), HL5136 rabbit anti-WHRN (Belyantseva et al., 2005), mouse anti-EPS8 (#610143, BD Biosciences), rabbit anti-GPSM2 (Ezan et al., 2013) and rabbit anti-GNAI3 (#G4040, Sigma-Aldrich). After washing in PBS, samples were labelled with Alexa Fluor 488 conjugated anti-IgG secondary antibodies (Life Technologies) diluted in blocking solution for 1 hour at RT. Samples were co-labelled with rhodamine phalloidin (Life Technologies) and mounted with high-precision #1.5 cover glass (Zeiss) using Prolong Gold (Life Technologies). Images were captured using a 63x objective (1.4 NA, plan apochromat, Zeiss) and a laser scanning confocal microscope (LSM780, Zeiss).

### Actin barbed end incorporation assay

Cochleae from *shaker-2* or *jordan* mice were dissected at P6 in Leibowitz L-15 media (Life Technologies) and adhered to the inside surface of a petri dish lid (Greiner CellStar 35 x 10 mm). The stria vascularis was left intact, as described (Xiong et al., 2014). Cochleae were cultured for 6 hours in DMEM / F-12 (Life Technologies) supplemented with 1% FBS (Atlanta Biologicals) and 1.5 µg·mL^-1^ ampicillin at 37°C, 5% CO_2_. Cochlea were incubated for 2 minutes at RT in 0.5 µM TMR-actin (Cytoskeleton) diluted in labelling buffer (concentrations in mM): HEPES (20), KCl (138), EGTA (3), MgCl_2_ (4), ATP (1), DTT (0.5), 1% (w/v) bovine serum albumin (BSA), 0.2 mg·mL^-1^ saponin, pH 7.5. Cochleae were immediately fixed for 30 mins at RT in 4% paraformaldehyde (EMS Diasum) diluted in PHEM buffer (concentrations in mM): PIPES (60), HEPES (25), EGTA (10), MgCl_2_ (2), sucrose (120), pH 7.0. Fixed cochleae were permeabilized in 0.5% (v/v) Triton X-100 in PBS for 30 minutes at RT, labelled with Alexa 488 phalloidin (Life Technologies), and mounted using a high-precision #1.5 cover glass (Zeiss) with Prolong Gold (Life Technologies). Confocal microscopy images were captured as described above.

### Expression of EGFP-MYO15-2 in mammalian cells

The pEGFP-C2-Myo15-2 and pEGFP-C2-Myo15-2(sh2) plasmids expressing an N-terminal EGFP fusion with the mouse MYO15 isoform 2 coding sequence (NP_874357.2) were previously reported (Belyantseva et al., 2003). The pEGFP-C2-Myo15-2(jd) plasmid was generated using site directed mutagenesis (QuikChange II, Agilent) to introduce the jordan (c.4940A>G) non-synonymous substitution. All expression plasmids were verified by Sanger sequencing and prepared as endotoxin-free, transfection grade DNA (NucleoBond Xtra Maxi EF, TakaraBio). HeLa cells (#CCL_2_) were obtained as an authenticated, low passage stock from the American Type Culture Collection (ATCC). LLC-PK1-CL4 (CL4) cells were a kind gift from Dr. James Bartles at Northwestern University. HeLa and CL4 cells were cultured in high-glucose DMEM (#11995, Life Technologies), supplemented with 1x GlutaMAX (Life Technologies) and 10% FBS (Atlanta Biologicals), and maintained at 37°C, 5% CO_2_. Transfection of HeLa cells or CL4 cells was performed using Lipofectamine 3000 (Life Technologies) according to the manufacturer’s protocol. pEGFP-C2 (Clontech) was used as an empty vector control. Cells were plated onto fibronectin (10 µg·mL^-1^) coated glass bottom culture dishes (#1.5, MatTek Corp) and allowed to adhere, and in the case of CL4 cells, characteristic island formation could be observed. Cells were fixed in 4% paraformaldehyde and 2% sucrose in PBS for 10 mins (EMS Diasum). Fixed cells were permeabilized/blocked in 0.2% (v/v) Triton X-100, 10% (v/v) normal goat serum (NGS) in PBS for 1 hr at RT, followed by labelling with rhodamine phalloidin (Life Technologies) and Hoechst 33342 (Life Technologies) in blocking buffer (2% NGS in PBS). Confocal microscopy images were captured as described above.

### Actin purification and labelling

Actin was extracted from rabbit skeletal acetone powder (Pel-Freeze, AZ) in chilled G-buffer (concentrations in mM), Tris-HCl (2), ATP (0.2), CaCl_2_ (0.1), NaN_3_ (1), DTT (1), pH 8, using established protocols (Spudich and Watt, 1971). Actin was additionally labelled on Cys 374 using either N-(1-pyrene)-iodoacetamide (Life Technologies), or tetramethylrhodamine-5-maleimide (Adipogen Life Sciences)(Criddle et al., 1985; Fujiwara et al., 2002). F-actin used for steady-state ATPase and gliding filaments assays was purified through two rounds of polymerization and depolymerization using ultracentrifugation. F-actin was dialyzed extensively against MOPS (4), MgCl_2_ (1), EGTA (0.1), DTT (1), NaN_3_ (1), pH 7.0, and its concentration measured at 290 nm (ε = 26,600 M^-1^·cm^-1^) prior to use. Unlabelled, pyrene- and TMR-labelled G-actin for polymerization studies were further purified by size exclusion chromatography (Superdex 200, Cytiva) using isocratic elution in G-buffer. Fractions were exclusively taken from the falling edge of the chromatogram, to ensure the recovery of monomeric actin. Concentrations and dye-labelling efficiency was determined at 290 nm (actin: ε = 26,600 M^-1^·cm^-1^), 344 nm (pyrene: ε = 22,000 M^-1^·cm^-1^) and 550 nm (rhodamine: ε = 96,900 M^-1^·cm^-1^). Correction factors were applied for pyrene actin, A_corr_ = A_290_ – (0.127 * A_344_), and separately for rhodamine actin, A_corr_ = A_290_ – (0.208 * A_550_)(Fujiwara et al., 2002). Biotinylated skeletal muscle actin (#8109-01, HyperMol, Germany) was rehydrated, dialyzed against G-buffer and cleared by ultracentrifugation for 60 minutes at 100k x *g* prior to use. For actin polymerization experiments performed without free ATP in solution, G-actin stocks (with 0.2 mM ATP) were desalted (PD SpinTrap G-25, Cytiva) into a modified G-buffer (no ATP): Tris-HCl (2), CaCl_2_ (0.1), NaN_3_ (1), DTT (1), pH 8, immediately prior to use. Desalted G-actin (ATP) monomers were stored on ice and used within 3 hours.

### Expression of MYO15 and deafness mutants in *Sf*9 cells using baculovirus

The baculoviral transfer vector pFastbac1 M15-2IQ-EGFP-FLAG, encoding the wild-type mouse MYO15 motor domain (NP_874357.2, aa. 1 - 743) as a C-terminal fusion with EGFP and FLAG moieties, was previously described (Bird et al., 2014). The expressed protein was 114 kDa. The *jordan* and *shaker-2* non-synonymous substitutions were separately introduced into pFastbac1 M15-2IQ-EGFP-FLAG by site-directed mutagenesis (QuikChange II, Agilent) and verified by Sanger sequencing. Plasmid DNA encoding wild-type, *jordan* and *shaker-2* was separately transformed into DH10Bac cells (Life Technologies) and recombinant bacmid DNA prepared following the manufacturer’s protocol. First passage (P1) recombinant baculovirus was generated by transfecting *Sf*9 cells (Expression Systems) with bacmid DNA complexed using linear polyethylenimine (PEI MAX, 40,000 MW, Polysciences Inc., PA) at a 12:1 (PEI : DNA) molar ratio. *Sf*9 cells were maintained in suspension culture with ESF-921 medium (Expression Systems) in a shaking incubator at 27°C. P1 baculovirus was amplified in *Sf*9 cells using low multiplicity of infection (MOI = 0.1) to generate P2 baculovirus for expression. Dual-promoter baculovirus expressing mouse UNC45B + HSP90AA1 chaperones (Bird et al., 2014), and bovine smooth muscle essential (MYL6, also referred to as MLC17B / ELC) + chicken regulatory (MYL12B, also referred to as MLC20 / RLC) light chains, were previously described (Pato et al., 1996). All baculoviruses were tittered using an end-point dilution assay and the *Sf*9 Easy Titer cell line (Hopkins and Esposito, 2009). To express M15-2IQ, *Sf*9 cells were seeded at a density of 2 x 10^6^ cells·mL^-1^ in ESF-921, and infected simultaneously with three baculoviruses at the following MOI: M15-2IQ (5), UNC45B / HSP90AA1 (5), ELC / RLC (5).The myosin chaperones UNC45B and HSP90AA1 we co-expressed to aid folding, in addition to essential (MYL6) and regulatory (MYL12B) muscle light chains to bind the LCBDs (Bird et al., 2014; Jiang et al., 2021). Identical expressions were performed for the *shaker-2* and *jordan* variants. Cells were harvested at 48 - 72 hours post-infection by centrifugation at 500 x *g* and flash frozen in liquid nitrogen.

### Purification of the MYO15 motor domain

M15-2IQ (wild-type, *jordan*, *shaker-2*) motor domains were purified from frozen *Sf*9 cells following established protocols (Bird et al., 2014; Jiang et al., 2021). Cell pellets were lysed using a Dounce homogenizer in extraction buffer (concentrations in mM): MOPS (10), NaCl (500), EGTA (1), MgCl_2_(10), ATP (2), PMSF (0.2), DTT (0.1), NaN_3_ (1), leupeptin (2 μg·mL^−1^), protease inhibitor cocktail (Halt EDTA-free; Thermo Scientific), pH 7.2. Cell lysates were cleared for 30 mins at 48k x *g* and the supernatant incubated with FLAG M2 affinity resin (Sigma-Aldrich) for 3 hours at 4°C. FLAG resin was packed into a gravity flow column and washed with a high-salt buffer, MOPS (10), NaCl (500), EGTA (1), MgCl_2_(5), ATP (1), PMSF (0.1), DTT (0.1), NaN_3_ (1), leupeptin (2 μg·mL^−1^), pH 7.2, followed by a low-salt buffer, MOPS (10), NaCl (60), EGTA (1), PMSF (0.1), DTT (0.1), NaN_3_ (1), leupeptin (2 μg·mL^−1^), pH 7.0. M15-2IQ protein was eluted using low-salt buffer supplemented with 0.2 mg·mL^−1^ 3x FLAG peptide (American Peptide, CA). For assessment of hydrodynamic radius, purified motor domains (wild-type, *jordan*, *shaker-2*) were concentrated by centrifugation (10’000 MWCO; Amicon, EMD-Millipore) and directly analyzed using size exclusion chromatography (see below), alongside protein size standards (Thyroglobulin (1), Ferritin (2), Aldolase (3), Conalbumin (4), Ovalbumin (5), Carbonic anhydrase (6), Ribonuclease A (7); HMW + LMW gel filtration calibration kit (Cytiva). For preparative scale protein production, FLAG-eluted motor domains (wild-type + *jordan*) were bound to a strong anion exchanger (5/50 MonoQ GL; Cytiva) using a Purifier 10 chromatography system (GE Healthcare). The column was washed with MOPS (10), NaCl (100), EGTA (1), PMSF (0.1), DTT (1), pH 7.0, and a 160 CV gradient elution performed to 1M NaCl (100% B). Fractions eluting at ∼31 mS·cm^−1^ were concentrated (10,000 MWCO) and further purified using size exclusion chromatography (Superdex 200, Cytiva) with isocratic elution in MOPS (10), KCl (100), EGTA (0.1), NaN_3_ (1), PMSF (0.1), DTT (1), leupeptin (1 μg·mL^−1^), pH 7.0. The M15-2IQ : ELC : RLC complex (1:1:1) eluted as a single peak and was concentrated (10,000 MWCO) before determining complex concentration at 280 nm (ε = 88,020 M^−1^·cm^−1^).

### ATPase assays

Steady-state actin activated ATPase assays were measured using a NADH-coupled assay, as previously reported (Bird et al., 2014). Briefly, wild-type M15-2IQ (30 nM) or *jordan* (150 nM) protein was assayed in the following reaction buffer (concentration in mM): MOPS (10), KCl (50), MgCl_2_ (5), EGTA (0.1), MgATP (2), 40 U·mL^-1^ lactate dehydrogenase (Sigma-Aldrich), 200 U·mL^-1^ pyruvate kinase (Sigma-Aldrich), phosphoenolpyruvate (1) (Sigma-Aldrich), NADH (0.2), pH 7.0 at 20° ± 0.1 °C. Due to its lower activity, a higher concentration (150 nM) of M15-2IQ *jordan* protein was used. The concentration of F-actin in the reaction was titrated from 0 to 100 μM. The absorbance of NADH (ε = 6,220 M^-1^·cm^-1^) at 340 nm was measured using a dual-beam spectrophotometer (UV-1800, Shimadzu) and the ATP hydrolysis rate calculated from the change in absorbance over time. Hydrolysis rates were corrected for basal M15-2IQ ATPase activity in the absence of F-actin, and also for the ATPase activity of F-actin. ATP hydrolysis rates were fit to the Michaelis-Menten equation to estimate *k*_cat_ and *k*_ATPase_ using Prism (GraphPad).

### Gliding filament motility assay

Motility chambers were assembled by coating a clean cover glass (#1.5) with 0.1% nitrocellulose in amyl acetate (Ladd Research Industries), and attaching it to a microscope slide with two strips of double-sided adhesive tape (Scotch, 3M) to form a channel approximately 3 mm wide. The chamber was incubated for 5 mins in 0.1 mg·mL^−1^ anti-GFP (clone GFP-20, Sigma-Aldrich) diluted with motility buffer (MB) (concentrations in mM): MOPS (20), KCl (10), MgCl_2_ (5), EGTA (0.1), pH 7.0. The surface was blocked using 1 mg·mL^−1^ BSA (Sigma-Aldrich) diluted in MB and incubated for 1 min. After washing the chamber with MB, 1 μM M15-2IQ (wild-type or *jordan*) diluted in MB was incubated for 1 min to functionalize the anti-GFP coated surface. The chamber was washed with 1 mg·mL^−1^ BSA in MB, followed by MB alone. Finally, the chamber was incubated for 2 mins with TMR-phalloidin stabilized actin filaments (5 nM) diluted in MB, and subsequently washed in MB. Filaments were visualized using an inverted epifluorescence microscope (Olympus IX-51) and motility recorded in MB supplemented with 50 mM DTT, 2 mM ATP, 3 mM glucose, 100 μg·mL glucose oxidase, and 20 μg·mL catalase at 30 ± 0.5 °C. Actin filament velocities were analyzed as described (Sellers et al., 1993).

### Polymerization assay

Actin polymerization was measured using G-actin labelled on Cys 374 with N-(1-pyrene)-iodoacetamide (see above), and a cuvette-based fluorometer (PTI Quantamaster 400, HORIBA Scientific) used to excite pyrene at 365 nm and capture fluorescence emission at 407 nm. Gel filtered G-actin (10% pyrene labelled, with free ATP, or desalted) was converted to the physiological Mg^2+^ bound form by addition of 50 µM MgCl_2_ and 0.2 mM EGTA for exactly two minutes at room temperature. The polymerization reaction was initiated by mixing G-actin (3 x stock) with KMEI buffer (1.5x stock) in a 1:2 ratio, respectively. Myosin (wild-type or *jordan*) was included in the 1.5x KMEI buffer as needed. Final concentrations were 2 µM G-actin and 1 µM myosin, in assay buffer (concentrations in mM): KCl (50), MgCl_2_ (1), EGTA (1), imidazole (10), pH 7.0 at 25° ± 0.1 °C. Data were corrected for dead-time and fluorescence recorded until the transient reached plateau, or for a maximum of 3 hours. Fluorescence derivatives with respect to time were processed in Prism (GraphPad). Half times were calculated as described (Hansen et al., 2013).

### TIRF single-filament polymerization assay

High-tolerance cover slips (24 × 50 mm, #1.5, Marienfeld Superior, Germany) were cleaned by sequential sonication (10 mins each) in 2% Hellmanex III (Hellma, Germany), 1M KOH, 100% ethanol, and finally Milli-Q water. Cover slips were dried under a nitrogen stream and processed for 10 mins under argon plasma (ZEPTO, Diener Electronic, Germany). A mixture of mPEG-silane (2 mg·mL^-1^, Laysan Bio, AL) and biotin–PEG–silane (2 μg·mL^-1^, Laysan Bio) was prepared in 96% ethanol, 0.1% (v/v) HCl. Cover slips were coated with 100 μL of the mPEG mixture and baked at 70°C for 1 hour. Cover slips were rinsed twice in 96% ethanol, sonicated for 10 mins, followed by two rinses in Milli-Q, sonicated for 10 mins in Milli-Q, and finally dried under a nitrogen stream. Flow chambers were assembled using double-sided sticky tape to create a 3 mm wide channel on a glass slide.

Functionalized coverslips were placed over the tape and firmly pushed down to seal the flow chamber. Immediately prior to use, flow cells were washed with buffer T50 (concentrations in mM): Tris·HCl (10), KCl (50), DTT (1), pH 8.0. Flow cells were sequentially washed with 0.1 mg·mL^-1^ (Neutravidin, Thermo Scientific) in T50 for 1 min, followed by a wash with 1 mg·mL^-1^ bovine serum albumin (A0281, Sigma Aldrich) in T50 for 30 s, and a final wash of T50. Experiments were performed in the following reaction buffer (final concentration in mM): KCl (50), MgCl_2_ (1), EGTA (1), imidazole (10), ATP (0.025), DTT (10), glucose (15), 0.5% methylcellulose, 20 μg·mL catalase, 100 μg·mL glucose oxidase, pH 7.0. G-actin (1 µM total) was included in the reaction with TMR-actin (20%) and biotin-actin (10%) doping. Purified myosin motor domain (1 µM) was optionally added, as needed. The reaction was loaded into the flow cell and immediately mounted onto an inverted microscope (Nikon Ti-E2) equipped with an oil immersion objective (CFI Apochromat TIRF 100x, 1.49 N.A., Nikon) for objective-style total internal reflection fluorescence (TIRF) microscopy (H-TIRF, Nikon). The flow cell was illuminated using a 561 nm laser line, and emission filtered using a bandpass filter (ET630/75m, Chroma). Time-lapse images were captured on an EM-CCD camera (iXon Ultra 888, Andor) controlled by NIS-Elements (AR version 5.2, Nikon). Samples were imaged at 21 ± 1°C. The assay dead-time was typically 45 - 60 s, and was included in data analyses.

Images were pre-processed in FIJI (https://fiji.sc)(Schindelin et al., 2012) by performing background subtraction and image registration (descriptor-based series registration, 2d/3d +t). Actin filament densities were quantified using the Analyze Particle command (size > 3 pixel^2^, circularity: 0.0-1.0) to count discrete particles within a 50 x 50 μm region of interest (ROI) that was randomly selected from the field of view. A minimum of 3 experiments, from two independent protein preparations, were analyzed for each condition. Filament elongation rates were calculated from time-lapse TIRF imaging sequences using kymographs generated in Elements Software (version 5.2, Nikon). At least 40 filaments from two independent experiments were analyzed for each condition.

### Statistical analyses

All statistical calculations and non-linear regressions were performed in Prism (GraphPad). Regression fit parameters are quoted as mean ± SEM, and data points as mean ± SD, unless otherwise stated. Statistical significance: * p < 0.05, ** p < 0.01, *** p < 0.001, **** p < 0.0001.

## AUTHOR CONTRIBUTIONS

Conceptualization: JEB, MRB

Investigation: ZGM, FJ, CA, MB, CFA, AS, JW, RH, DCS, SMC, AP, SM, EW, YT, MRB, JEB

Formal analysis: ZGM, FJ, CA, MB, RG, AS, CFA, JW, RH, DCS, SMC, AP, SM, EW, YT, JRS, SDMB, GMA TBF, MRB, JEB

Writing - original draft: ZGM, FJ, CA, MB, MRB, JEB Writing – review and editing: All authors.

Project administration: JEB

Funding acquisition: JRS, SDMB, TBF, MRB, JEB

## ACKNOWLEDGEMENTS

We thank Mireille Montcouquiol for the gift of GPSM2 antisera, James Bartles for the gift of LLC-PK1-CL4 cells, Tracy Fitzgerald at the NIDCD Mouse Auditory Testing Core for expert technical assistance, W. Clay Smith for use of fluorometry equipment, Peter Höök, H. Lee Sweeney and Tom Pollard for insightful discussions, and Peter Barr-Gillespie for critical feedback on the manuscript. This research was supported (in part) by the Intramural Research Program of the NIH (NIDCD DC000039 to TBF, NHLBI HL006049 to JRS), the Medical Research Council (MC_U142684175 to SDMB, MC_UP_1503/2 to MRB) and National Institute on Deafness and Other Communication Disorders of the National Institutes of Health under Award Number R01DC018827 (to JEB). The content is solely the responsibility of the authors and does not necessarily represent the official views of the National Institutes of Health. The funding bodies had no role in study design, data collection or analysis, decision to publish, or preparation of the manuscript.

## ACKNOWLEDGEMENTS

The authors declare that no competing interests exist.

---

**FIGURE S1.**
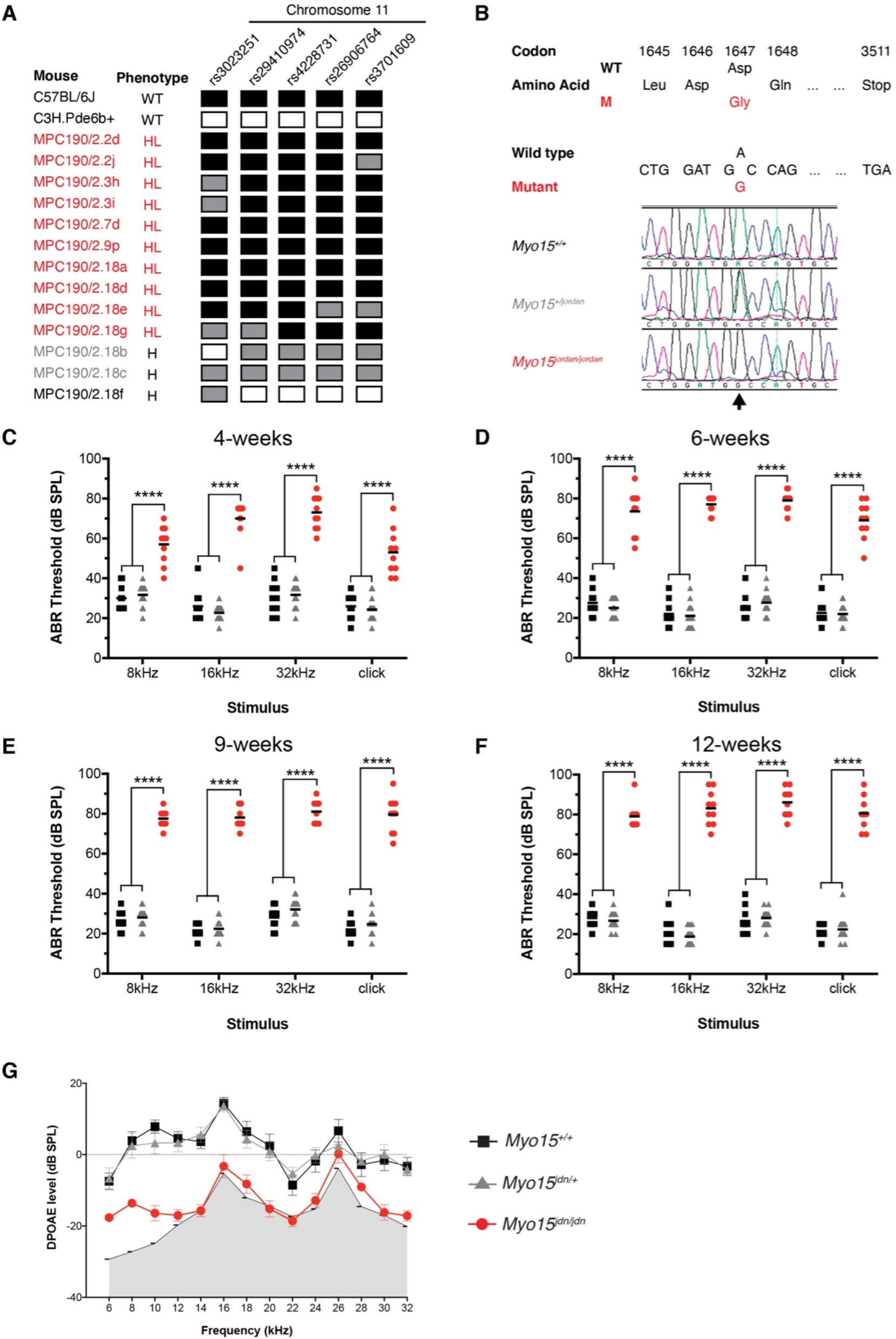
**A)** Whole genome SNP mapping of genomic DNA from ten MPC190 mice exhibiting hearing loss (red, HL) and three hearing (H) littermate controls. The genotype of each mouse is either homozygous for C57BL/6J (black box), C3H (white box) or heterozygous (grey box) for each marker. The analysis defined a 16.5Mb critical interval on Chromosome 11 between markers rs29410974 and rs26906764 (Chr11:50420012-67162951, GRCm38). **B)** Sanger sequencing confirmed the only WGS-identified, coding lesion (arrow) within the critical interval, in exon 17 (ENSMUSE00000244718) of the *Myo15* gene (ENSMUSG00000042678). A nucleotide transition (c.4940A>G) at codon 1647 alters the wild-type aspartate (D) codon to a mutant glycine (G). Example electropherograms are shown for *Myo15^+/+^*, *Myo15^+/jd^* and *Myo15^jd/jd^*. **C-F)** Minimum ABR detection thresholds for a longitudinal cohort of *Myo15^jd/jd^* mice (n = 10), *Myo15^+/+^* (n=10) and *Myo15^+/jd^* (n=15) littermates at 4, 6, 9 and 12 weeks reveal progressive hearing loss. Data points represent individual mice, with the mean value indicated. **** p < 0.0001; two-way ANOVA with Tukey’s multiple comparisons test. **G)** Distortion Product Otoacoustic Emission (DPOAE) responses are greatly reduced at 12 weeks in *Myo15^jd/jd^* mutants (red, n = 10) at all frequencies tested compared with *Myo15^+/+^* (black, n = 9) and *Myo15^+/jd^* (gray, n = 14) littermates. Data are mean ± SD.

**FIGURE S2.**
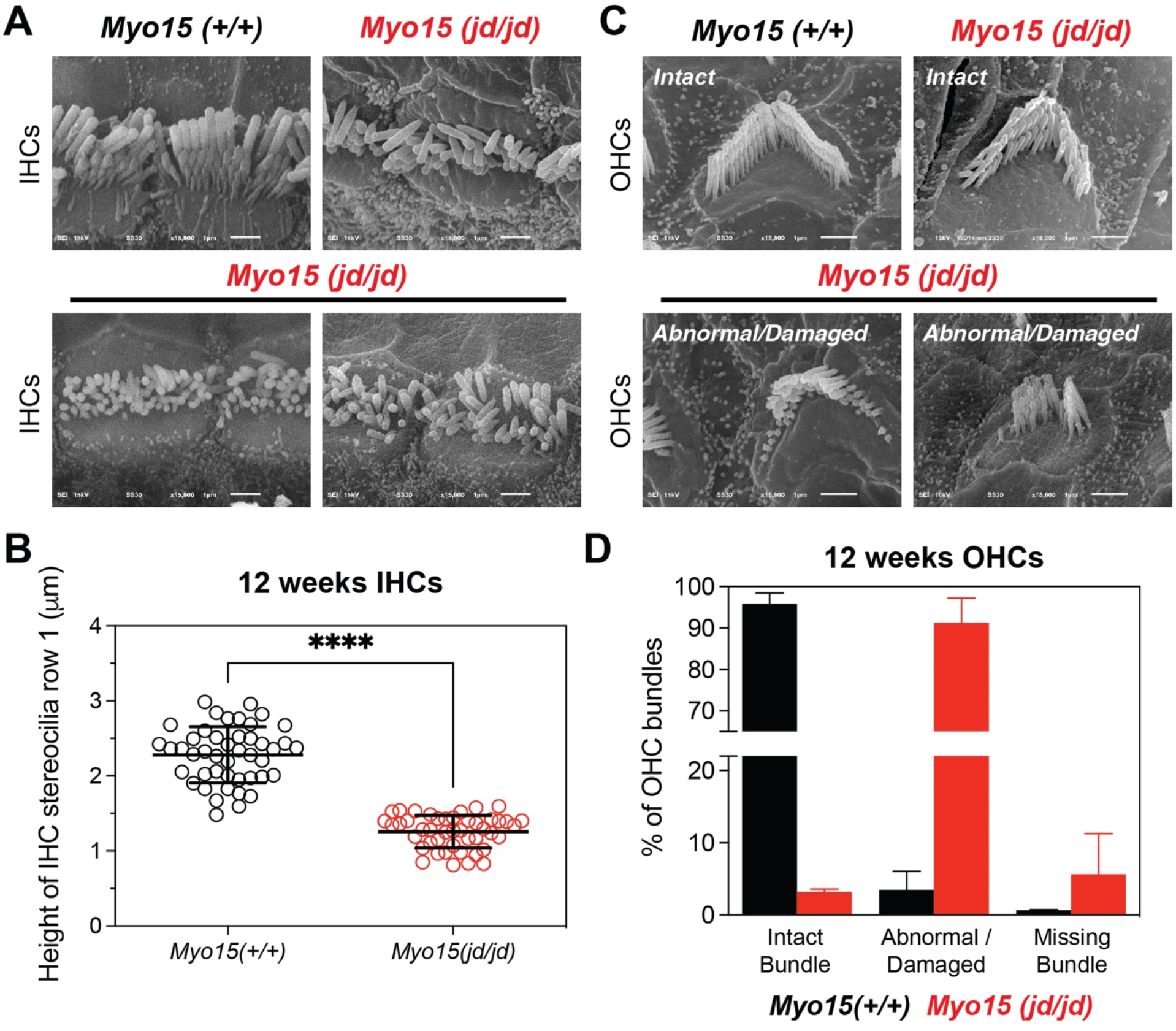
**A)** SEM analyses of IHC bundles from *Myo15^jd/jd^* and *Myo15^+/+^* littermates at 12 weeks. **B)** Quantification of the tallest (row 1) stereocilia at 12 weeks in IHCs from *Myo15^+/+^* mice (45 stereocilia from 3 animals) and *Myo15^jd/jd^* mice (45 stereocilia from 3 animals). **** *p* < 0.0001, unpaired, two-tailed *t*-test. **C)** SEM analyses of *Myo15^jd/jd^* OHC bundles at 12 weeks. OHC bundles were qualitatively categorized as “intact”, “abnormal/damaged” or “missing”. Representative examples are shown. In the “abnormal/damaged” group, stereocilia were typically resorbed from the periphery of the hair bundle. **D)** Quantification of OHC bundle categories at 12 weeks in *Myo15^+/+^* (1344 OHCs from 4 animals) and *Myo15^jd/jd^* littermates (972 OHCs from 4 animals). All images and quantification are from the mid-cochlear turn. Scale bars, 1 µm.

**FIGURE S3.**
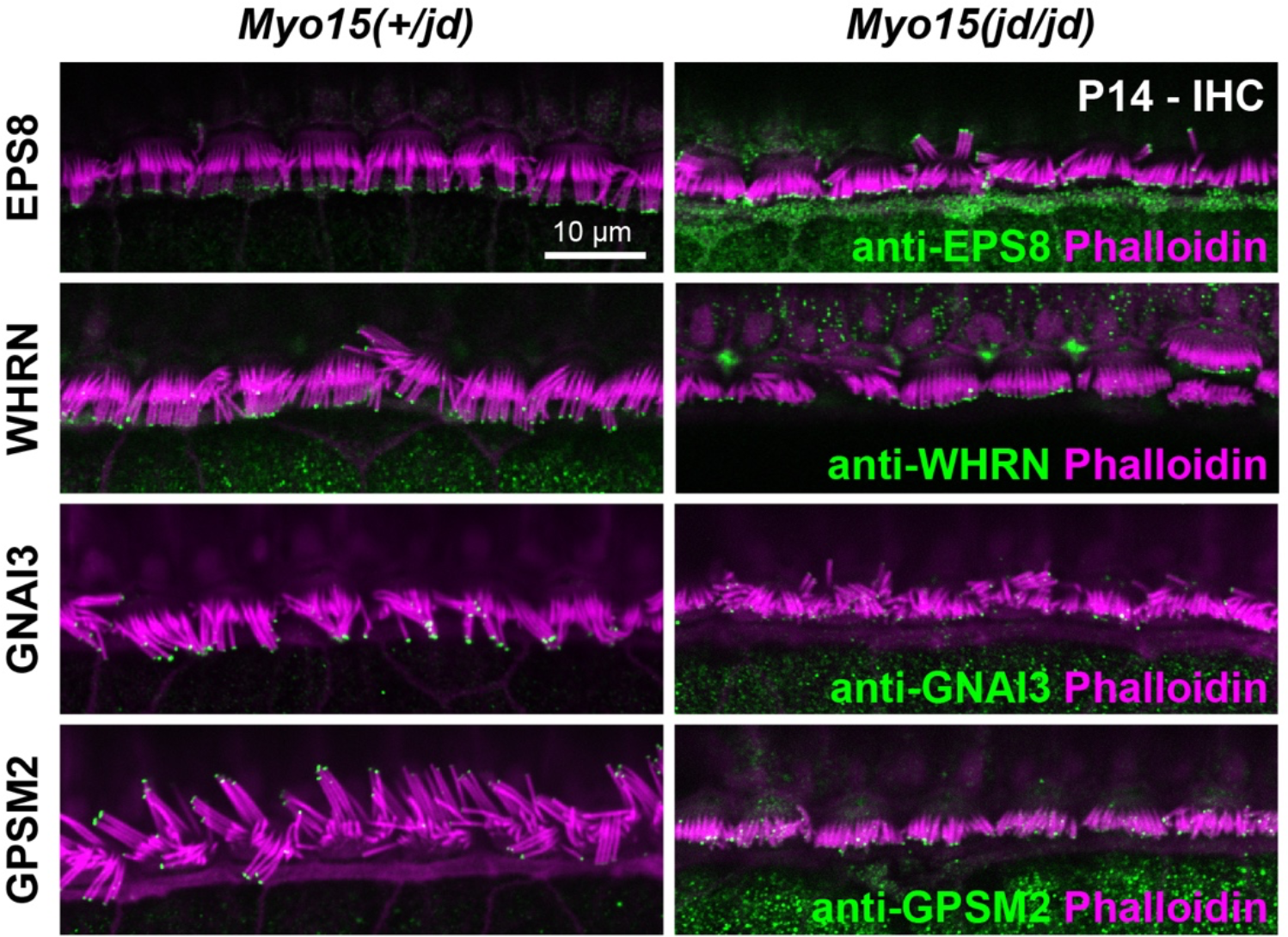
Immunofluorescence (IF) confocal images of anti-EPS8, anti-WHRN, anti-GNAI3 labelling or anti-GPSM2 (green) in control *Myo15^+/jd^* and *Myo15^jd/jd^* IHCs fixed at P14. Phalloidin was used to label F-actin (magenta). Images are representative of data from two independent animals per genotype and antibody combination. Scale bars, 10 µm.

